# Social and Spatial Affinity Drive Wound Care by Ants

**DOI:** 10.1101/2025.10.08.681151

**Authors:** Ebi Antony George, Alba Motes-Rodrigo, Laurent Keller, Erik Frank

## Abstract

In group-living animals, open wounds pose a risk not only for the injured individual but also for the entire group, especially when the risk of infection transmission is high. To mitigate this danger, different species have developed diverse strategies – with ants representing one of the few taxa beyond humans that engage in social wound care. Yet, who provides this care and what determines its allocation remains unclear. To answer these questions, we combined controlled injury experiments with automated behavioral tracking across six colonies of the ant *Camponotus fellah*. Our results reveal that wound care is primarily provided by a non-specialized subset of individuals whose behavioral traits align with a transitional phase between the nurse and forager roles. Among these transitioning ants, care provision was best predicted by their prior spatial and social proximity to the injured individuals in the hours before the injury, indicating that pre-existing social affinity influences caregiving behavior. These findings offer the first empirical evidence of how caregivers are integrated into the colony’s social structure and highlight the role of social interactions in shaping the organization of wound care in ants.

## Main text

Open wounds in animals can arise in various contexts, such as competition for resources or predation, and can severely compromise individual fitness by increasing the risk of mortality. To mitigate this threat, many species employ self-directed wound care behaviors that promote healing and reduce the risk of infection such as wound licking to remove debris, applying antimicrobial compounds^1,2^, and smearing pharmacologically active plants as topical self-medication^3,4^. In contrast to these self-care behaviors, social wound care, a cornerstone of human healthcare systems, has only been documented in a few taxa including primates^5–7^, ungulates^8^, rodents^9^ and eusocial insects. In ants, social wound care involves behaviors like wound cleaning^10,11^, the application of antimicrobial substances^12^ and even amputation of injured limbs by nestmates^13,14^. These behaviors form part of a broader phenomenon known as social immunity, a suite of collective strategies employed by insect colonies to reduce infection risk and maintain colony integrity^15^.

Previous research on social immunity has primarily focused either on group-level responses by mapping, for example, changes in social network topology^16^ or on individual behavioral performance by characterizing grooming or wound care behaviours^10^. However, the critical link between these two organizational levels – specifically how the colony-level social structure shapes and regulates individual caregiving – remains largely unexplored.

One possibility is that wound care is carried out by a dedicated subset of specialists, akin to nurses or foragers who consistently perform brood care or foraging tasks, respectively^17^. Alternatively, caregiving might reflect a transient behavioral state adopted by non-specialist individuals predisposed to engage in care due to specific traits, prior interactions with the injured individual, or both. Such temporary predispositions to engage in tasks have been observed in other social species. For example, in the ant *Myrmica rubra*, the tendency to remove sporulating corpses correlates with individual levels of prophylactic self-grooming^18^. Similarly, in ungulates and primates, prior affiliative interactions between individuals can predict helping behaviors^19–21^. Yet, in the context of social immunity, we lack a detailed understanding of how individual characteristics and interaction history influence the propensity to provide care.

To investigate whether wound care is provided by a specialized caste, and to understand the role of caregivers within the colony’s community structure as well as the factors driving care allocation, we conducted a large-scale behavioral study. We combined high-resolution automated tracking of 660 ants from six *Camponotus fellah* queenright colonies (Fig. 1B) with a targeted injury experiment. In this experiment, foragers were identified based on their proximity to food sources and injured in a sterile environment by severing the right hind leg at the femur. Following injury, each ant was returned to its colony, and its interactions were manually annotated for six hours post manipulation (450 hours of video data). We annotated three behaviors: wound care (i.e., cleaning the injured leg with the mouthparts), allogrooming, and trophallaxis (Table S1). Allogrooming is a well-established component of social immunity^22^, whereas trophallaxis has rarely been investigated in this context. However, trophallaxis may play an important role by facilitating information transfer regarding health status among nestmates or by providing nutritional compensation during immune responses^23^. To complement our behavioral data, we used automated tracking to continuously record the trajectories and social interactions of all ants in each colony (1440 hours of tracking data and 787,986 interactions across colonies), enabling detailed spatial and social profiling of caregivers before and after injury events.

**Figure 1:**
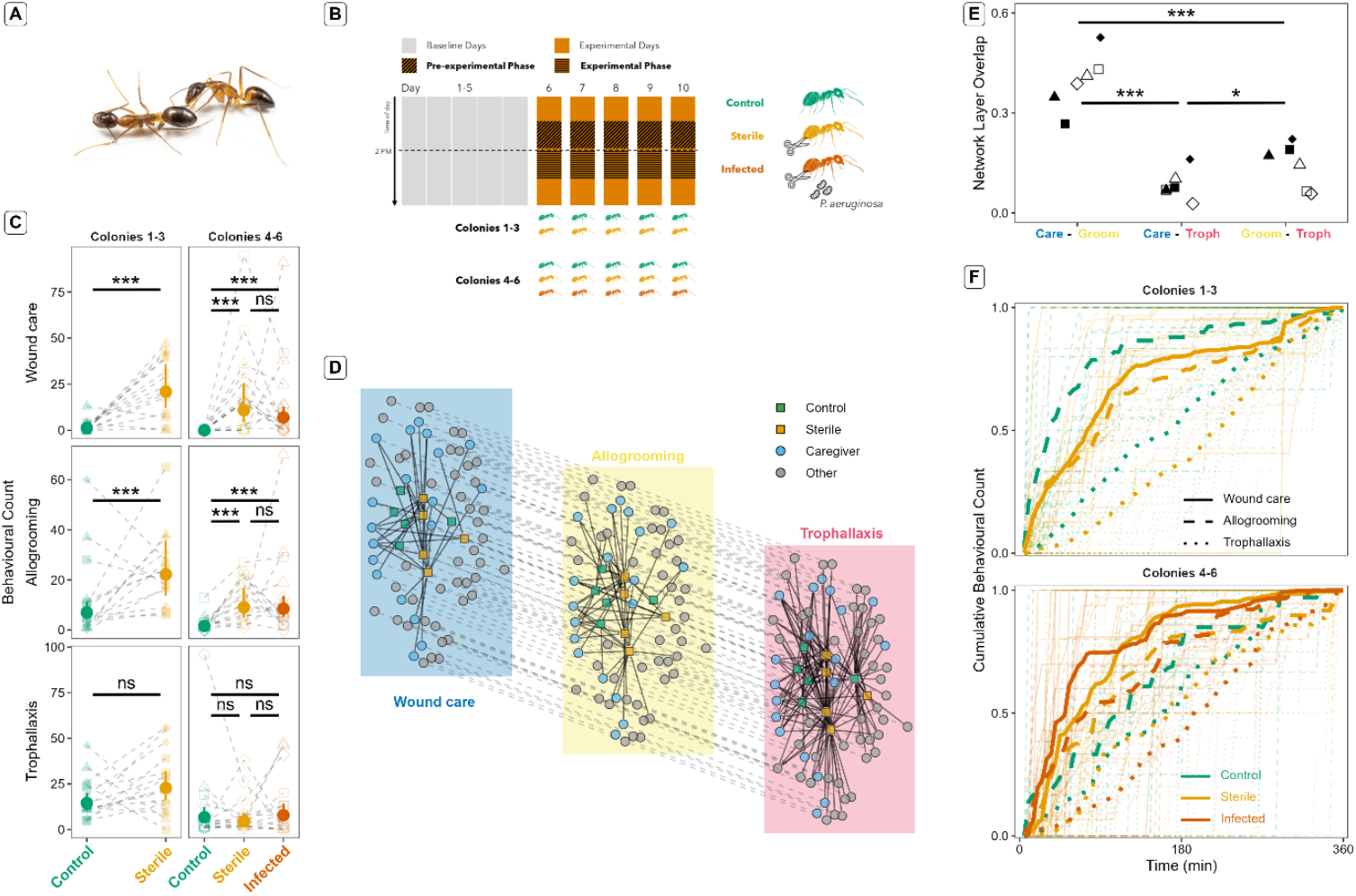
A) An injured ant receiving wound care. We observed a total of 89 individual caregivers from six colonies (14.83 ± 4.62 caregivers per colony) and approximately 4.67 ± 2.07% of ants provided care on any given day per colony. In total, we recorded 982 wound care events (17.85 ± 21.17 events per injured ant). B) Experimental protocol for each replicate colony and focal group. Experiments were conducted over five consecutive “Experimental days” (Day 6 – 10) following a five day “Baseline” period where colonies acclimatized to the tracking setup (Days 1 – 5). On each experimental day, behavioral data were analyzed for six hours immediately post-manipulation (“Experimental phase”). Spatial and interaction data were collected for both the Experimental phase and the six hours preceding the manipulation (“Pre-experimental phase”). Focal groups are represented by the names next to the ants and different colors: sterile = ants in orange with sterile injuries, infected = ants in red with infected injuries and control = uninjured ants in green. C) Frequency of the target behaviors during the Experimental phase. Solid circles and vertical lines represent the estimated marginal means and 95% confidence intervals of the number of occurrences for wound care (cleaning of the target leg), allogrooming and trophallaxis. Semi-transparent points represent data from individual ants with filled shapes indicating colonies 1-3 and open shapes indicating colonies 4-6. Different colors denote each focal group (as in B). Dashed lines connect focal ants from the same experimental day in each colony. D) Representative egocentric multiplex behavioral network. The network represents the interactions of nestmates with focal ants (egocentric around the focal ants) with each layer corresponding to a specific behavior (multiplex). It is based on data from all five experimental days of Colony 1 and includes all interactions with five control ants and five ants with sterile injuries. Nodes represent individual ants, and the edges are weighted by the duration of the behavioral interaction between pairs of ants (weighted equally for visualization). Grey dashed lines across layers connect the same ant in the different layers. E) Network layer overlap between pairs of behavioral layers. Different shapes denote data from individual colonies (filled shapes: colonies 1-3, open shapes: colonies 4-6). F) Cumulative distribution functions of the timings of events for all three target behaviors. Opaque lines represent the average of the distribution function for each focal ant type and semi-transparent lines represent the distribution for individual focal ants. The color scheme is the same as in B. Significance indicators: ns for p > 0.05, * for 0.01 < p < 0.05, *** for p < 0.001.

To verify that sterile wounds elicit wound care and to test whether trophallaxis and allogrooming frequencies increase post-injury, we performed controlled experiments using three colonies (colonies 1 to 3). Each day for five consecutive days, we selected two foragers per colony and randomly assigned them to either a control (uninjured) or a sterile injury group. We hypothesized that injured ants would receive more wound care, allogrooming and trophallaxis than controls. As predicted, injured ants received significantly more wound care and allogrooming (Fig 1C, Tables S2-S4). However, the amount of trophallaxis was not higher for injured ants than controls (Fig 1C, Tables S2-S4).

In *Megaponera analis*, wound care is modulated by infection status, with infected wounds receiving more frequent treatment^12^. Conversely, studies in *Camponotus maculatus* found no such difference between sterile and infected injuries^14^. To assess whether *C. fellah* responds differently to infected wounds, we conducted a second experiment using colonies 4–6. Each day for five days, we randomly assigned three foragers per colony to one of three groups: uninjured, sterile injury, or infected injury. Infected ants had their right hind leg severed, and the injury exposed to a *Pseudomonas aeruginosa* solution in phosphate-buffered saline. A preliminary survival experiment confirmed that infected wounds reduced survival compared to sterile injuries (Fig. S1, S2), and we hypothesized that infected ants would receive more care.

Contrary to our expectations, the frequency of wound care and allogrooming did not differ significantly between sterile and infected groups (Fig 1C; wound care – difference = 0.45, z = 0.82, p = 0.96; allogrooming – difference = 0.07, z = 0.18, p = 0.86). Furthermore, the presence or absence of infected ants had no effect on the treatment received by ants with sterile injuries (colonies 4-6 vs 1-3; Table S3 and S4; difference in wound care received per ant = 1.07, z = 2.05, p = 0.56). Taken together, these findings parallel those observed in *C. maculatus*^24^, suggesting that, unlike *M. analis*^10^, wound care in *Camponotus* species such as *C. fellah* may be prophylactic rather than dependent on the wound’s infection status.

Our behavioral observations also revealed a distinct separation between care provisioning (allogrooming and wound care) from food provisioning of injured ants in terms of which individuals performed these behaviors and when they were performed (Fig 1D-F). An analysis of egocentric multiplex networks (Fig. 1D) showed that wound care and allogrooming were performed by a smaller and more overlapping set of individuals compared to trophallaxis. Specifically, the network overlap between wound care and allogrooming was significantly higher (wound care network nodes = 20 ± 4.69, allogrooming = 35 ± 6; overlap between networks = 0.40 ± 0.09) than between either behavior and trophallaxis (trophallaxis nodes = 55.70 ± 14.70; overlap between wound care and trophallaxis = 0.08 ± 0.04, allogrooming and trophallaxis = 0.14 ± 0.07). Furthermore, nestmates which spent more time caring for the wounds of injured ants also groomed them more but there was no correlation between time spent on wound care and trophallaxis (Fig. S3, Pearson’s correlation coefficient: wound care-allogrooming = 0.52, p < 0.001; wound care-trophallaxis = -0.03, p = 0.50).

The temporal patterns of these three behaviors also differed markedly. The majority of wound care and allogrooming events (approximately 75%) occurred within the first two hours post-injury and drastically decreased in frequency thereafter (Fig 1F). In contrast, trophallactic interactions involving injured ants occurred at a consistent rate throughout the six hour observation period (Fig 1F, Table S8 and S9; difference in median time of occurrence of wound care vs trophallaxis for ants with sterile injuries in colonies 1 to 3 = -1.04, t = -4.85, p < 0.001; colonies 4 to 6 = -0.59, t = -2.12, p = 0.075; for infected ants = -1.07, t = -3.77, p = 0.001). Trophallactic networks ensure the circulation of a variety of substances within the colony and play a vital role in maintaining colony cohesion^25,26^. Our results show that these networks are structurally and temporally independent of care provision networks.

To characterize the drivers of wound care, we first assessed whether simply encountering an injured ant was sufficient to elicit care. We compared the temporal distribution of encounters between nestmates and injured ants with the distribution of wound care events, hypothesizing that if merely encountering an injured ant led to care, both would follow similar patterns. However, unlike wound care, which occurred immediately post-injury (Fig. 1F), nestmates encountered the injured ants consistently throughout the six-hour observation period (Fig. S4, difference in median time of occurrence of wound care vs encounters = -62.94, t = -4.19, p < 0.001). This temporal mismatch indicated that encounters alone were insufficient to explain the observed wound care dynamics.

We next tested the hypothesis that only the timing of encounters determines whether wound care takes place, with earlier encounters being more likely to lead to care. Alternatively, nestmate identity could be important, and nestmates could have different probabilities of providing care regardless of their probability of encountering injured ants. To differentiate between these hypotheses, we used the time-ordered encounter data of all nestmates with the injured ants (n = 30 injured ants) to simulate wound care provisioning under the condition that every ant had the same probability of providing care (only the timing of encounters determines whether care takes place). Our aim was to see if these simulations could reproduce both the observed temporal dynamics of wound care and the actual number of caregivers. Conversely, if nestmates differed in their probabilities of becoming caregivers, then these simulations would not fully replicate our observations. We ran simulations using either constant probabilities of care applied uniformly across all encounters and injured ants (probabilities of Slow: 0.05, Smed: 0.25, Shigh: 0.5) or an empirically derived probability for each injured ant which remained constant for all encounters with that specific ant (Sempirical probability for an injured ant = the ratio of wound care events received to the number of encounters; mean probability across injured ants = 0.08 ± 0.11, range = 0.007 - 0.45).

Our simulations revealed that even the scenario that best approximated the observed temporal dynamics of wound care (Slow; Fig. 2A; for a comparison of observed and expected median times per ant see Table S10), failed to reproduce the empirically observed number of caregivers (Fig. 2B). In fact, the actual number of caregivers was consistently and significantly lower than predicted across all simulation scenarios (Fig. 2B; Table S10). Therefore, encounter dynamics alone, assuming equal care probabilities amongst nestmates, were not sufficient to reproduce the observed wound care patterns. Instead, these results strongly support our alternate hypothesis that not all nestmates are equally likely to provide care and that a small group of individuals are predisposed to wound care.

**Figure 2:**
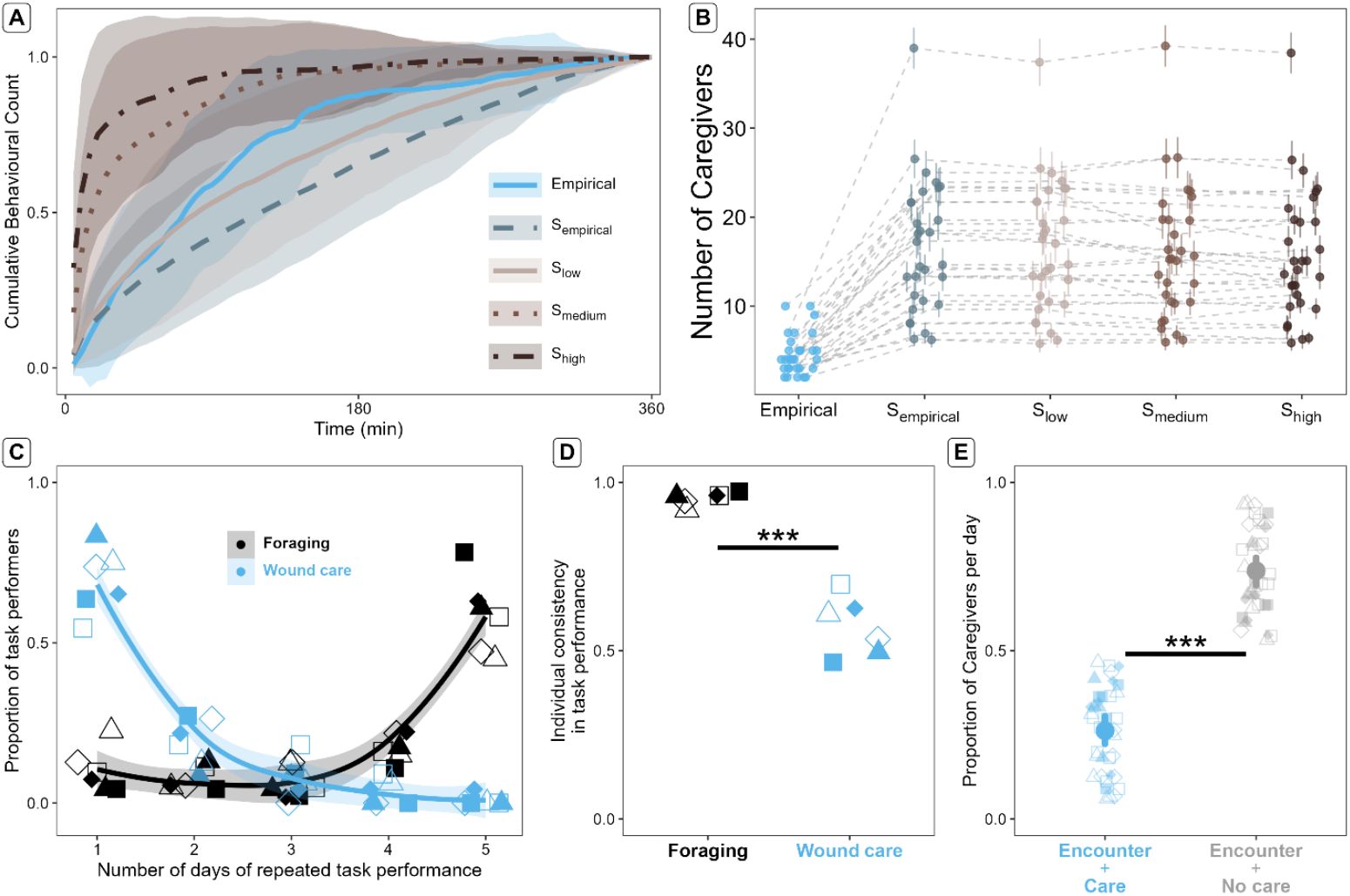
A) Cumulative distribution of wound care events. The average cumulative distribution of wound care events received by injured ants (n = 30, including ants with sterile and infected wounds) from empirical observations (blue solid line) and four simulation scenarios (S_empirical_, S_low_, S_med_, S_high_; represented by gray and brown lines of different types). The shaded region represents the standard deviation around the mean. B) Observed versus simulated number of caregivers. Each point represents the number of caregivers for one injured individual. Grey dashed lines connect data from the same individual across observed and simulated conditions. The circles and error bars represent the mean and standard deviation from 1000 simulations. The color scheme is the same as in A. C) The proportion of workers engaged in wound care and foraging repeatedly over days. Each point represents the proportion of all identified caregivers (blue) or foragers (black) who performed their respective tasks on multiple days per colony (filled shapes: colonies 1-3, open shapes: colonies 4-6). The solid line represents a LOESS curve fit to the data from all colonies, with the shaded region indicating the confidence interval around this line. D) Individual consistency in foraging and wound care. Individual consistency in a given task is measured using a modified mutual entropy-based division of labor statistic which accounts for repetition in task performance over days. Each point represents data from one colony with the color denoting the task and the shape denoting the colony as in C. E) The proportion of caregivers who encountered the injured ant and either provided care or did not. Solid circles and vertical lines represent the estimated marginal means and 95% confidence intervals of the proportion. Semi-transparent points represent data from individual days with different colors for the two groups of caregivers and different shapes for each colony as in C. Significance indicators: *** for p < 0.001.

To assess whether caregivers constitute a specialized group, we compared their consistency in task engagement with that of foragers. As expected, foragers showed a high degree of consistent engagement in their task with most individuals repeatedly foraging on all experimental days in each colony (proportion of foragers active on all 5 days per colony = 0.59 ± 0.12). In contrast, most ants that engaged in wound care only did so on one day (Fig. 2C; mean ± standard deviation = 1.46 ± 0.85 days, median = 1 day). Moreover, only a small proportion of caregivers (mean proportion per colony = 0.20 ± 0.19) provided care to both sterile injured and infected injured ants even when both ants were present in the colony at the same time. Quantifying this consistency in repeated task performance using a standard mutual entropy-based division of labor statistic^27^ (see Supplementary material), we found that caregivers had a significantly lower consistency than foragers (Fig. 2D; difference = -2.67, z = -10.95, p < 0.001). This lack of repeated care was not due to a lack of encounters between caregivers and injured ants, as 74% of all caregivers who encountered injured ants did not provide care on a given day (Fig. 2E, difference = -2.07, z = -13.29, p < 0.001). Taken together, these results show that, contrary to foraging, wound care is not consistently performed by a specialized group of workers.

In *C. fellah*, a prominent division of labor among nestmates leads to two separate communities of workers performing different tasks and occupying distinct nest areas^28^. Nurses are typically younger individuals which engage in intranidal tasks like brood care and as they age, they transition into the forager community. Ants transitioning between the two communities overlap in their space use and interactions with members of both communities^29^. To evaluate where caregivers fit within this well-defined community structure, we compared their spatial and social characteristics with those of nurses and foragers, using established community-defining spatial and social metrics (time spent inside the nest, activity, nest coverage and interactions with members from the other communities, for definitions of communities see supplementary material).

Our results revealed that ants that engaged in wound care showed spatial and social characteristics typical of individuals transitioning between the nurse and forager communities^28,30^. Caregivers spent significantly less time in the nest than nurses, but significantly more time than foragers (Fig. 3A and 3B; caregivers vs nurses: difference = -1.88, z = -13.49, p < 0.001; caregivers vs foragers: difference = 0.49, z = 3.58, p = 0.001). They were also significantly more active than both nurses (t = 9.41, p < 0.001) and foragers (t = 2.52, p = 0.042; Fig. S4 and Table S10) and covered a greater portion of the nest than both groups (Fig. 3B; p < 0.05 for both comparisons). This extensive nest coverage by caregivers resulted in caregivers being on average 7.82 times more likely to interact with nurses and 27.1 times more likely to interact with foragers than expected by chance based on their group sizes (difference estimate between the two normalized interaction rates = 16.72, t value = 1.87, p = 0.067). Reciprocally, nurses and foragers were on average 1.35 and 5.75 times more likely to interact with caregivers, and this was significantly higher than their interaction rates with the opposite community (nurses with foragers = 0.40 ± 0.30, difference estimate = 0.96, t value = 3.82, p < 0.001; foragers with nurses = 0.70 ± 0.60, difference estimate = 5.00, t value = 10.12, p < 0.001). In *C. fellah*, individual workers take about 20 days to transition from the nurse to forager communities, and at any given time, around 20% of the colony’s workers are in this process^29^. Our findings indicate that wound care is performed by ants in this critical developmental stage.

**Figure 3:**
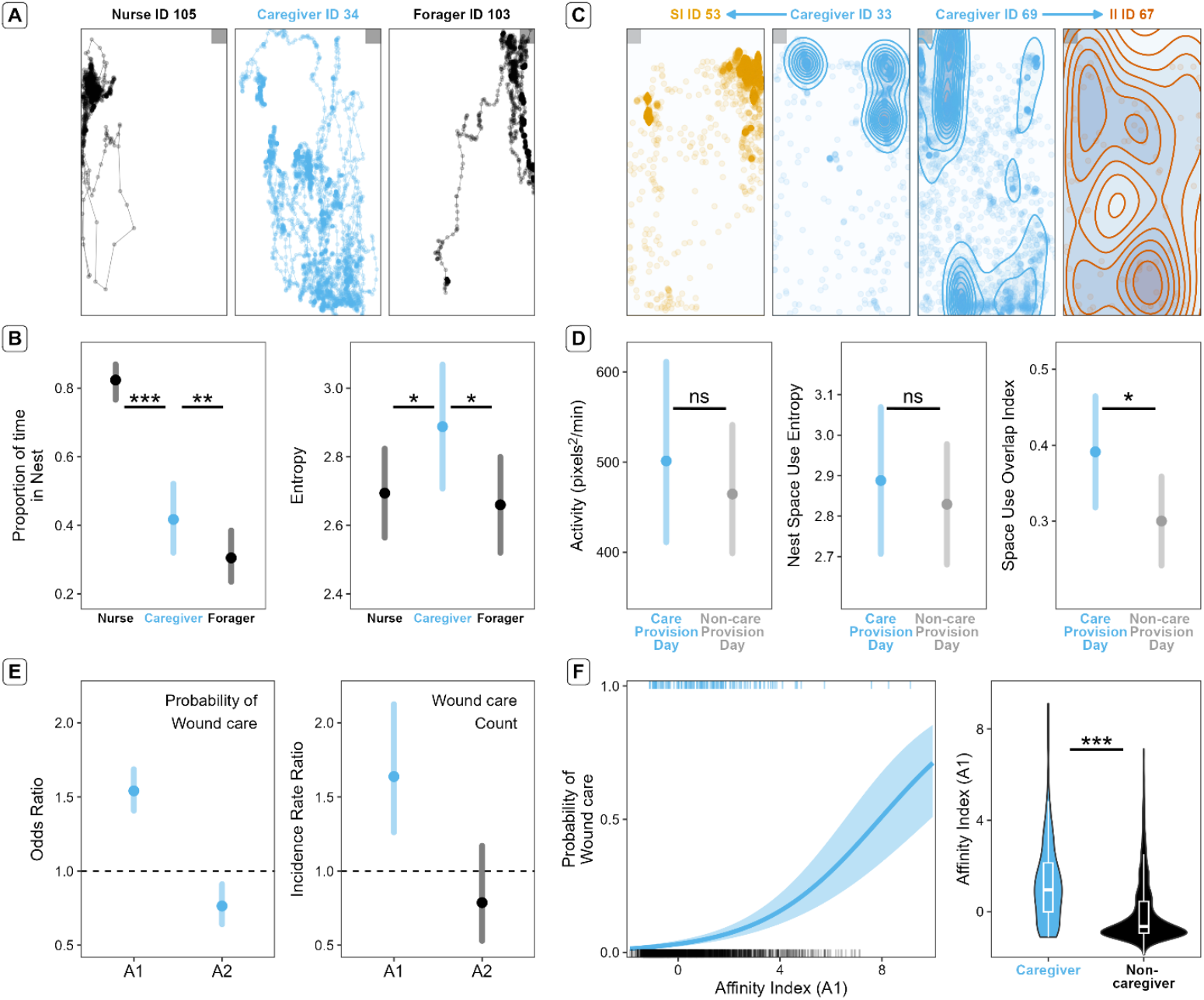
A) Representative trajectories of ants from different task categories. Trajectories in the nest arena of a nurse (identified by its social maturity value), a forager (identified by its presence in the foraging arena), and a caregiver during the 2 hours before manipulation on experimental day 2 in colony 2. The grey boxes indicate the nest entrance. Black lines represent data for nurses and foragers and blue for caregivers. B) Comparison of spatial characteristics between task categories. The proportion of time spent in the nest and the nest space use entropy of nurses, caregivers and foragers. Solid circles and vertical lines represent the marginal means and the 95% confidence intervals. The color scheme is the same as in A. C) Representative kernel density estimates of space use of focal ants and caregivers. The space use in the foraging arena during the six hours before manipulation for an ant with a sterile injury (ID 53), two caregivers (ID 33 and ID 69), and an infected ant (ID 67) on experimental day 4 from colony 5. The colors of the kernels correspond to the ant type with orange for the ant with a sterile injury, red for an infected injured ant and blue for caregivers. The grey boxes indicate the exit towards the nest. Blue arrows above the trajectory panels indicate the specific injured ant each caregiver provided care to in the Experimental phase. D) Comparison of caregiver characteristics on days on which they provided care versus days on which they did not. The activity, nest space use entropy and space use overlap are compared between the two groups. Solid circles and vertical lines represent the marginal means and the 95% confidence intervals. Blue represents data from care provision days and grey from non-care provision days. E) The effect of prior affinity on wound care provision. The effect of A1 and A2 (the first two principal components obtained from the affinity related properties) obtained from the Pre-experimental phase on the probability (odds ratio) and the amount (incidence rate ratio) of wound care provided after the injury. Solid circles and vertical lines represent the effect size and 95% confidence limits. The colors represent the statistical significance of the effect size (blue for significant, grey for non-significant). The dashed line at the effect size of one indicates the threshold at which a change in the predictor has no effect on the probability or amount of care. F) Predicted increase in probability of wound care with change in affinity. The blue line describes the estimated probability of an ant becoming a caregiver in the Experimental phase as a function of its affinity index (A1) value in the Pre-experimental phase. The shaded region represents the confidence intervals around this estimate (x-axis has been zoomed in for visualization). Rug plots inset on the top and bottom show the distribution of the affinity index values for individuals which provided care (blue) and those which did not (black). The violin plots show the distribution of A1 for caregivers and non-caregivers during the Pre-experimental phase. Significance indicators: ns for p > 0.05, * for 0.01 < p < 0.05, *** for p < 0.001.

However, being in a transitional stage between communities is not sufficient to fully explain care provision, as individuals only provided care on intermittent days and to specific injured ants (Fig. 2E, Fig. S4). Furthermore, if being in this transitional stage solely explained care, then caregivers should show significant differences in intrinsic behavioral traits that characterize transitioning ants between days they provided care and days they did not. Instead, caregivers did not significantly differ in their spatial and social characteristics on care provision days compared to non-care provision days (Fig. 3C, 3D, S4 and Table S11). Thus, caregivers’ intrinsic spatial and social characteristics are not sufficient to explain our observations on wound care behavior.

We then tested whether a caregiver’s spatial and social affiliation with injured ants prior to the injury was crucial to their decision to provide care. We hypothesized that if prior affiliation was important, then caregivers would show higher affiliation with the injured ants on days they provided care compared to days in which they did not. To test this, we quantified the overlap in space use and the number of interactions between caregivers and foragers six hours prior to the injury. In line with our expectations, we found that caregivers shared significantly more space with the injured ants on days they provided care compared to days they did not (Fig. 3C, 3D, Table S10; difference in overlap index = 0.09, t = 2.82, p = 0.018). However, the number of interactions with injured ants remained the same (Table S10, difference in interactions = 0.35, z =1.47, p = 0.39). Combined, these results indicate that while intrinsic traits associated to transitioning stages may predispose ants to provide care, selective care provision is directly linked to their pre-existing spatial affiliation with the injured ant.

To verify these findings, we assessed whether we could predict which individuals amongst all the nestmates in the colony were more likely to provide care on a given day based on their behavior prior to the manipulation. To do so, we quantified 13 different individual properties from three different behavioral domains (an individual’s spatial properties, its position in the colony’s social network, and its affinity with the soon-to-be injured ants) measured during the six hours before injury, along with the first two principal components of each property type for all ants in the colony (Table S12 for individual properties and loadings; see Supplementary Methods for further details). We then built and compared 12 statistical models using these properties, each testing a specific hypothesis, to identify the best predictors of care provision. The model incorporating A1 and A2, the first two principal components representing an ant’s prior affinity with the injured individual, had the highest predictive power (BIC = 1090.66, relative weight = 0.90), surpassing all other models including those with single variables like the space use overlap or the number of previous interactions with injured ants (relative weight of next best model = 0.05, Table S13).

A1, which we refer to as the “affinity index”, strongly correlated with greater space use overlap, more encounters and more interactions with the injured ant prior to the injury. This affinity index showed a positive relationship with the probability of care provision: a unit increase in affinity led to a 54% increase in the probability that an individual would provide care on a given day (baseline odds-ratio = 0.03, odds-ratio of A1 = 1.54; Fig 3E, 3F, Table S13). Moreover, unlike A2, which correlated with the number of connections needed to reach the injured ant in the interaction network, the affinity index A1 also predicted the overall amount of care provided, and not just the probability of providing it (Fig 3E, Table S14, incidence rate ratio of A1 = 1.64, z = 3.71, p < 0.001; A2 = 0.79, z = -1.18, p = 0.24). Collectively, these findings demonstrate that the likelihood and extent of an ant providing wound care are best predicted by its prior spatial and social affinity with the injured individual.

Overall, our study describes, with unprecedented resolution, who and what drives care provisioning in ants. We found that social care in *C. fellah* is not the result of ants merely encountering injured individuals. Instead, non-specialized workers who are in a transitional state between nurses and foragers are predisposed to engage in wound care. Crucially, an individual’s probability to engage in wound care on any given day is determined by its prior spatial and social proximity to the injured nestmate. Our findings offer novel insights into how the dynamic social and spatial organization of ant colonies enables social immunity strategies based not on specialized individuals but rather on the affinity between workers. Such affinity-driven care may represent a widespread, yet underappreciated, mechanism for maintaining group cohesion and resilience in social insects and other group-living organisms.

## Supporting information

Supplementary materials, methods and results

