## Supplementary materials, methods and results for "Social and Spatial Affinity Drive Wound Care by Ants"

* shared first authors
[+] shared last authors
[1] Department of Ecology and Evolution, University of Lausanne, Lausanne, Switzerland
[2] The Sense Innovation and Research Center, Lausanne and Sion, Switzerland
[3] Social Evolution Unit, Chesières, Switzerland
[4] Department of Animal Ecology and Tropical Biology, University of Würzburg, Würzburg, Germany

### Supplementary Materials

### Materials and Methods

#### Subjects and housing

We used six colonies of *Camponotus fellah* collected in 2007 (one colony) and 2010 (5 colonies) in Tel Aviv, Israel. At the time of the experimental manipulations the colonies had been in the laboratory between 13 and 16 years. Each colony contained between 200 and 500 workers, a reproductive queen and brood in all three stages of development (egg, pupae and larvae). Workers showed dimorphism into minors and majors. Colonies were housed following standard protocols[28,29] inside plastic containers with walls covered in Fluon, an extremely smooth substance that prevents ants from climbing out. The containers included drinking tubes filled with water and a nest structure. Ants were fed twice a week with honey water solution (1:5 parts honey:water) and frozen Drosophila flies. Ant colonies were kept in a room at 26ºC, 65% humidity and a 12h light cycle. We selected 110 workers per colony balancing ants picked from the nest and near the honey water containers to ensure a proportionate representation of foragers and nurses in the experimental subcolony. Approximately 10% of the selected workers were majors. The queen and approximately 60 brood items were also included in each experimental subcolony. Experimental subcolonies were formed 1h before being introduced into the behavioral tracking arena.

#### Automated behavioral tracking

Immediately after the subcolonies were formed, each ant was tagged using unique fiducial markers (ARTags) measuring 1.3 x 1.3 mm that were glued to the thorax of each ant using skin glue (Manfred Sauer GMBH). Ants were anesthetized on ice before tagging. Two tracking arenas were used in each colony replicate, one representing the nest and one representing a foraging arena. Both arenas were rectangular (22 x 17 x 9.5cm), had Fluon covered walls, a non-reflective foam floor and a drinking tube filled with water. The nest arena was kept in darkness for the duration of the study and maintained at a constant temperature of 26 ºC during the day and 25 ºC during the night with a humidity of 65% throughout. The foraging arena was set to cycles of 12 h light and 12 h darkness, a temperature of 26 ºC during the day and 25 ºC during the night with a humidity of 55% throughout. Each ant colony was tracked for 11 days in total (days 0 to 10). Ants were fed five flies and honey water at the beginning of the tracking period (day 0) in the foraging arena and on each experimental day during the Experimental phase (days 6 to 10, Fig 1B). To allow the ants to acclimatize to the new environment, the tracking data for the first day, when ants were introduced into the system, was discarded (day 0).

Behavioral tracking was done using the FORmicidae Tracker described in[31]. Briefly, this system consists of a monochromatic camera (45 megapixel with a 40 mm focal length wide angle lens) paired with an infrared illumination system. The system captures images at 5Hz and stores the spatial coordinates of each tag, along with video recordings of the arenas (https://formicidae-tracker.github.io). The experiment in each colony lasted five days and was conducted immediately after the baseline tracking period (baseline days: 1 - 5; experimental days: 6 - 10). The baseline days tracking data was used to calculate the baseline activity and different behavioral measures of all ants in an undisturbed condition as described in the statistical analyzes below.

### Experimental manipulations

In three of the colonies (1 to 3), two tagged minor workers were selected each day from the foraging arena and randomly assigned to a focal group (control or sterile injured). Ants were selected from the foraging arena because these ants had a higher probability to be foragers and foragers are the most likely ants to experience injuries in the wild[10]. All ants were anesthetized on ice before applying the manipulation. Sterile injured ants received a cut on the right hind leg approximately at the center of the femur with sterile scissors[12]. The control ants were exposed to the disinfected scissors in the same location as injured ants but without injuring it. Immediately after, ants were returned to the foraging arena. This was repeated with a new pair of ants on five consecutive days, for a total of five injured and five control ants per colony (total = 15 ants per focal group across colonies).

In the other three colonies (4 to 6), three tagged minor workers were selected each day from the foraging arena and randomly assigned to a focal group (control, sterile injured or infected injured). Control and sterile injured ants were treated with the same protocol as described above for colonies 1 to 3. For the injured infected ants, we infected the wounds using *Pseudomonas aeruginosa*, a common gram-negative bacteria found in the soil[32], to which ants are regularly exposed in the wild. The pathogen was isolated from soil samples from Côte d'Ivoire [12]. To ensure that the bacteria used to perform the wound infections were in the same growth phase in each of the five experimental days, the bacteria was plated from frozen stock into TSA anew 48h before each experimental day. Bacterial dilutions and infections followed the protocol described in the "Test of *Pseudomonas aeruginosa* concentration lethality" below. After the manipulation, ants were returned to the foraging arena. In total, we had five sterile injured ants, five infected injured ants and five uninjured ants as control per colony (total = 15 ants per group across colonies 4 to 6).

To decide on the bacterial concentration to use to infect the ants, we performed a 6-day survival experiment to determine a suitable bacterial concentration that would reduce individual survival compared to a sterile injury if no wound treatment was provided.

#### Test of *Pseudomonas aeruginosa* concentration lethality

We compared the life-expectancy of ten uninjured ants (control), ten ants injured with sterile wounds and 30 ants injured with wounds infected with one of three concentrations of *P. aeruginosa* (Optical density (OD): 0.0005, 0.005, 0.01, ten ants per concentration). Control ants were exposed to the sterilized scissors used to perform the injuries but no wound was inflicted and no solution was applied to the leg. The wounds of the ants in the sterile injured group were dipped into a 10 µl drop of sterile PBS for three seconds. Infections took place by dipping the ant's injured leg into a 10 µl drop of bacterial solution for three seconds. All infections took place on the same day from a stock stored in tryptic soy broth (TSB) with 25% glycerol at -23 °C. Two days before the infections, bacteria were plated in tryptic soy agar (TSA) medium and stored at room temperature for 24h. Isolated colonies were then transferred to liquid TSB medium and stored at room temperature for 24h. Thirty minutes before the manipulation of the ants, phosphate-buffered saline (PBS) solutions with a given OD were prepared from the liquid stock and the bacterial concentrations measured (using a Novospec III visible spectrophotometer, Amersham Biosciences). All materials were sterilized between manipulations with 90% ethanol. Ants in the survival experiment were continuously filmed while in isolation inside sterile petri dishes with a feeder containing honey water. All ants tested were minor workers and they were considered dead when they had not moved or twitched for more than 30 minutes while lying on their side. Based on the results of this experiment (Fig. S1), we used a bacterial concentration of 0.005 OD in the main manipulations, as done in other wound infection studies on ants [12,13,24].

#### Infection survival experiment

In parallel to the main manipulation in which infected injured workers were re-introduced into their colonies of origin (colonies 4-6), we conducted a second survival experiment to ensure there was no cross-contamination between the sterile and infected injuries. This also allowed us to additionally check that the plated pathogens were still active and affected individual mortality. Each experimental day, three foragers (minor workers) were selected from the colony of origin (not the sub-colony, to maintain group size of the tracked individuals) and assigned to each of three manipulation groups – ants with no injuries, with a sterile injury or and infected injury with the 0.005 OD dilutions prepared on that day. Manipulations in the three groups were applied following the same protocols as described above. These ants were continuously filmed while in isolation in individual sterile petri dishes with a feeder filled with honey-water. All ants tested in both survival experiments were considered dead when they had not moved or twitched for more than 30 minutes while lying on their side (Fig. S2).

### Behavioral coding and automated quantification of interactions

Each experimental day, we manually coded the behavioral interactions of the focal ants (control, sterile injured and infected injured) during the six hours following the manipulation (Experimental phase), as most wound care events have been previously described to occur in this time window[10–13,24]. For each interaction with the focal ants, we recorded the group of the focal ant (i.e., control, sterile injured or infected injured), the identity of the other ant involved in the interaction, the timestamp at the start of the interaction, the timestamp at the end of the interaction, the arena in which it took place (nest or foraging arena) and the type of interaction. The definition of the different behavioral interactions can be found in the ethogram in Table S1. The behaviors were annotated by two observers with high inter-rater reliability (based on a comparison of 50 randomly selected events, Cohen's Kappa = 0.975). In total, we manually annotated 450 hours of focal interactions (six hours per day, five experimental days per colony, six colonies, with two focal ants on each experimental day for colonies 1 to 3 and three focal ants on each experimental day for colonies 4 to 6).

Table S1: Ethogram. In the example images, blue ants represent focal ants (control, sterile injured or infected injured) and yellow ants represent caregivers or non-focal ants.

| Behavior | Depiction | Description | Example |
|---|---|---|---|
| Wound care | 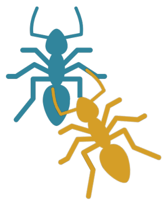 | Nestmate places mouthparts on injured ant at the location of the injury (on the third leg on the right side at the femur). | 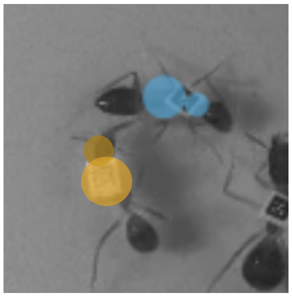 |
| Leg cleaning | 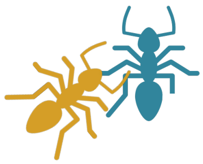 | Nestmate places mouthparts on legs of the focal ant. This is different from wound care as it does not take place at the wound and can be equated to allogrooming of the legs. | 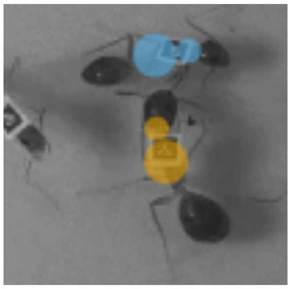 |
| Allogrooming | 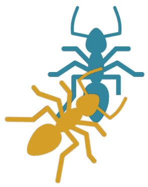 | Nestmate places its mouthparts on top or under the focal ants' body and moves its mandibles repeatedly over it. | 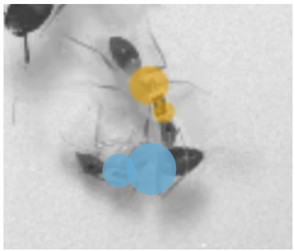 |
| Trophallaxis | 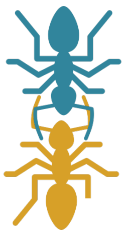 | Nestmate and the focal ant open and lock their mandibles in place, engaging in mouth-to-mouth fluid exchange. | 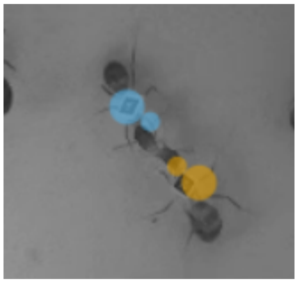 |

In addition to the manual behavioral annotation, we analyzed 1440 hours of tracking data per ant for all 660 ants. The tracking dataset was divided into the following phases: Baseline days (five days preceding the start of the manipulations); Baseline phase (period of six hours on the day preceding the first experimental manipulation in each colony); Pre-experimental phase (period of six hours before the manipulation); Experimental phase (period of six hours following the manipulation, starting at approximately 2pm each day) and Post-experimental phase (period of six hours, starting approximately 18 hours after the manipulation, Fig. 1B). For each of these phases we computed all nestmate interactions using the fort-studio post processing software[31]. To define proximity-based interactions, for each ant we drew a body shape consisting of two perpendicular ovals corresponding to the head and the abdomen. The body shapes were scaled to the individual ant size. An interaction started when the head oval of an ant overlapped with the head oval of another ant and ended when the head shapes no longer overlapped. Across the different phases and colonies, we quantified 787,986 interactions between pairs of ants. We also extracted the trajectories of all ants in the four 6-hour phases (Baseline, Pre-experimental, Experimental and Post-experimental) to look at their movement properties. This was not done for the Baseline days due to the large amount of data present over the five days which made it difficult to obtain the kernel density estimates for each ant's trajectories without downsampling the data.

#### Encounter-based classification of opportunities to provide care

Opportunities to engage in care were quantified as the number of encounters between a nestmate and an injured ant. An encounter was defined based on when an ant moved from beyond an outer threshold (distance between nestmate and injured ant, $\Delta d_n > D_O$) to within an encounter threshold around the injured ant ($\Delta d_n < D_E$). Based on the average size of individuals in our tracking recordings, we defined $D_E = 300$ pixels (~ 1 ant body length) and $D_O = 1000$ pixels (> 3 ant body lengths). For a new encounter to be considered, an ant had to move beyond $D_O$ and return to within $D_E$. This two-threshold measure allowed us to distinguish between instances where nestmates were still in the vicinity of the injured ant (beyond $D_E$ but within $D_O$) from new encounters when the nestmate and the injured ant came spatially closer together.

#### Statistical analyses

Models were fitted in R[33] with the functions lmer, glmer from the package lme4 and the function glmmTMB from the glmmTMB package[34,35]. We report the estimated marginal means and 95% confidence intervals around these means throughout the manuscript for the various statistical comparisons. Random slopes of factors were dummy coded before being included in models. Model assumptions, including collinearity and overdispersion, were checked using the check_model function from the performance package, the function VIF from the car package[36], by simulating residuals using the DHARMa package[37] or using diagnostic plots available in the stats package in R. Pairwise comparisons between factor levels were conducted using the function emmeans from the emmeans package[38] as well as using functions in the modelbased package[39]. The model structures for all the models are provided in Table S2.

For processing the tracking data, we used the py_myrmidon library to query the trajectory and interaction data (https://formicidae-tracker.github.io/myrmidon/latest/), pandas[40] and numpy packages[41] in python[42] for data handling and the networkx package in python for creating interaction edgelists[43]. Social maturity was calculated using facetnet[44] with a custom package (https://c4science.ch/source/facet_unil/).

For the general data analysis and plotting we used the tidyverse suite of packages[45] in R. We used igraph[46], ggraph[47] and MuxViz[48] in R for network analysis and the amt package[49] for space use calculations. For plotting, we used (in addition to ggplot2 from the tidyverse), ggh4x[50], cowplot[51], ggpp[52], patchwork[53], ggpubr[54] and graphlayouts[55]. From base R, we used the prcomp function for principal component analyses and cor.test for correlation analyses.

#### Comparison of care and food provisioning

*Behavioral response to focal ants*

We investigated differences in frequency (counts) and total duration (over the whole Experimental phase) of behaviors directed towards the focal ants. To evaluate whether leg cleaning was focused on the injury (and thus targeted towards the open wound indicating wound care), we also compared the duration and frequency of licking directed towards the other legs of the focal ants. Thus, for our comparison of wound care behavior we tested for the interaction effects between 'focal group' (control, sterile injured and infected injured) and 'leg' as a factor. The 'leg' factor consisted of two levels: treated leg and non-treated legs. This allowed us to test two specific hypotheses: a) whether injured individuals received overall more licking events, by comparing licking of the treated leg between the injured and control individuals and b) whether licking was specifically focused on the injured leg by comparing the licking received by the treated leg with that of the other five non-treated legs in the injured individuals. We fit six models evaluating whether the frequency and duration of leg licking (colonies 1 to 3: models M1, M2; colonies 4 to 6: M3, M4), allogrooming (colonies 1 to 3: M5, M6; colonies 4 to 6: M7, M8) and trophallaxis (colonies 1 to 3: M9, M10; colonies 4 to 6: M11, M12) differed between focal groups.

*Activity of focal ants*

To test whether behavioral differences between the focal ants were due to differences in their activity in the Experimental phase (colonies 1 to 3: M13; colonies 4 to 6: M14). We also checked whether the individuals we selected for the manipulations differed in their activity in the Baseline and Pre-experimental phase. We compared the average displacement (in pixels) of each focal ant per minute (displacement of the focal ants in each frame relative to the previous one aggregated per minute). We use the average instead of the total displacement as sometimes focal individuals may not be detected by the tracking system (e.g., when they are on their sides receiving care or moving in the tube connecting the foraging and the nest arena) and the average accounts for this missing tracking data.

*Social maturity of focal ants*

Social maturity is a metric ranging from 0 to 1 indicative of the integration of ants into the forager community computed using the facetnet community detection algorithm based on the social interaction network obtained from processing the tracking data[30]. Since all our focal ants were taken from the foraging arena, we expected them to be part of the forager community. However, they could vary in how embedded they are within the community and hence in their social maturity values. We compared the social maturity values of the focal groups to test the hypothesis that differences in the behavioral response of the nestmates to the focal ants are driven by how embedded the focal ants are within the forager community. We estimated the social maturity value of all focal individuals in each colony using data from the five baseline days. Differences in social maturity between focal ants were evaluated using a beta regression model (colonies 1 to 3: M15; colonies 4 to 6: M16).

*Behavioral multiplex network comparison*

For behavioral multiplex network comparisons and subsequent analysis, we considered wound care as cleaning of the target leg directed towards sterile and infected injured individuals. Cleaning of other legs of these individuals and cleaning of the target leg and other legs in control ants were considered allogrooming.

We evaluated the relationship between the individuals who performed the three behaviors by building edge colored multiplex networks. We combined data from all experimental days for each colony and created a three-layer network centered on the focal ants with each layer corresponding to a behavior, nodes corresponding to individual ants and edges weighted by the duration of the behavioral interaction between the nestmate and the focal ant. We first compared the size of these networks by quantifying the number of nodes in each layer (M17). Then we compared the network layer overlap between the layers to identify which layers were the most similar in their network connections and hence involved similar individuals (M18).

*Correlations between behaviors at the individual level*

To corroborate our findings at the colony level from the multiplex network analysis, we looked at correlations between wound care, allogrooming and trophallaxis at the individual level. For this, we obtained the total duration of all three behaviors performed by nestmates per injured ant and quantified the Pearson correlation coefficient between pairs of behaviors across all ants involved in the interactions with the injured ants.

*Behavioral dynamics over time*

To compare the time dynamics of behaviors, for each behavior (wound care, allogrooming and trophallaxis) we created a vector of time elapsed from the starting of the experimental phase (when the focal ants were reintroduced back into the foraging arena) until an event was observed (i.e., the relative latency of the event). We then generated an empirical cumulative distribution of these times, extracted the median value of the distribution and compared it across behaviors and focal groups. For colonies 1 to 3, we performed one comparison within injured ants to determine whether the median values of the latency distribution for wound care was different from the distributions for allogrooming and trophallaxis (M19). We performed another comparison including control and injured ants to test whether the median latencies for allogrooming and trophallaxis were different between them (M20). We performed the same comparisons for colonies 4 to 6 with the addition of the infected ants in both models (M21 and M22). To ensure that we had sufficient values to estimate the medians in each distribution, for each individual we only shortlisted those behaviors with at least five events.

#### Caregiver analyses

In this set of analyses, we investigated who provides care to injured ants in the colony. We first analyzed whether any nestmate which encounters injured ants provides care. We then focused on where caregivers lie within the existing division of labor structure in the colony. Next, we compared characteristics of caregivers with non-caregivers (individuals who have equal opportunities to provide care but do not do so) and differences within caregivers between days on which they provided care versus days on which they did not. Finally, we tested whether nestmates' characteristics before the injuries took place are linked to their propensity to provide wound care. Since the behavioral response of nestmates to the injured and infected ants were the same, we combined data from these two focal ant groups for this set of analyses.

*Encounter based simulations of wound care*

We first compared the temporal dynamics of wound care events with the encounters of nestmates with injured ants. The same process as described above in the section on "*Behavioral dynamics over time*" was used to extract the median value of the cumulative distributions of the latencies of both wound care events and encounters. We then statistically compared the median latencies between the two (M23). As before we shortlisted data from injured ants which had at least five care events and performed this comparison on 30 injured ants from all six colonies.

Next, we simulated different scenarios in which wound care is driven solely by the time-based encounters of nestmates with the injured ants and does not depend on individual differences in providing care. Our goal was to compare the temporal dynamics of care as well as the number of caregivers from the simulation of these scenarios with empirical data to determine if any of these scenarios were sufficient to explain our observations of wound care. We ran these simulations using the encounter data of all nestmates with injured ants during the 6-hour Experimental phase obtained from the automated tracking system. We simulated the wound care that each injured ant received (n = 30) using either a constant probability of wound care per encounter for all nestmates across all experiments (probabilities of $S_{low}$: 0.05, $S_{med}$: 0.25, $S_{high}$: 0.5) or an injured ant-dependent probability for all nestmates. For the latter scenario, we used the empirical data of each injured ant to calculate its probability of receiving care on any given encounter as the ratio of wound care events received to the number of encounters ($S_{empirical}$, with mean probability = 0.08 ± 0.11, range = 0.007 - 0.45). To conduct the simulations, we first ordered all the encounters of nestmates with the injured ants sequentially in time as empirically observed. We then simulated for each encounter whether wound care took place one after the other based on the probability that we defined for that scenario and injured ant. Each simulation run ended when the number of simulated wound care events was equal to the observed care events for that injured ant. In case the number of care events was not reached at the end of the encounters (e.g., in $S_{low}$), we then selected all the remaining encounters without care events, ordered them in time and re-started the sequential simulation of each encounter. This process maintained the time ordering of the encounters over multiple cycles and was repeated till we reached the observed number of care events. Thus, in these simulations, each encounter has a fixed probability of eliciting wound care and does not depend on whether the previous encounter elicited wound care. We simulated each of the four scenarios 1000 times per ant after shortlisting ants which received at least five wound care events (120,000 simulations in total from 30 ants). Our comparisons between $S_{low}$, $S_{med}$, $S_{high}$, $S_{empirical}$ and the empirical data allowed us to understand whether simulating wound care as a response of nestmates all of whom had the same fixed probability explains the wound care dynamics we observed.

We obtained two outputs from each simulation run: which encounter led to a wound care event and the ID of the nestmate which provided this care. Using the time of the simulated encounter which elicited care, we formed an empirical cumulative distribution of simulated wound care and extracted the median time as described in the previous analysis. For each ant in each scenario, we obtained 1000 median times and used this to create a cumulative distribution of median times per ant as our null distribution. We then compared how extreme the value of the observed median time of wound care was under this null distribution. For the number of caregivers, we performed a similar analysis. We first obtained the total number of caregivers from each simulation for each ant, created a null distribution using the 1000 values per ant for each scenario and compared how extreme the value of the observed number of caregivers was under this null distribution.

#### Consistency in task performance of wound care

In social insects, several tasks like foraging and nursing, are performed by a specialized group of workers. These workers repeatedly engage in this task over multiple days. We tested whether caregivers are a as specialized as foragers by quantifying the consistency in individual care behavior over the five days for each colony and comparing this consistency with that of foragers. We defined caregivers as any individual which provided wound care at least once and foragers as any individual present in the foraging arena at least once during the same Experimental phase during the five experimental days. To quantify consistency within each task, we used a modified form of a mutual entropy-based division of labor statistic (DOL). For each task we calculated DOL as:

$$DOL = \sqrt{DOL_{indiv} \times DOL_{day}}$$

where

$$DOL_{indiv} = \frac{I\,(indiv, day)}{H\,(indiv)}$$

and

$$DOL_{day} = \frac{I\,(indiv, day)}{H\,(day)}$$

Here $I(indiv, day)$ is the mutual entropy over the joint distribution of task performance by individuals over the five experimental days and is given by

$$I\,(indiv, day) = H\,(indiv) - H\,(indiv|day) = H\,(day) - H\,(day|indiv)$$

*H (indiv)* and *H (day)* represent the standard Shannon's index of entropy over individual and daily variation in task performance respectively, given by

$$H(X) = -\sum_{x \in X} p(x) \times \log\,[p(x)]$$

and *H (indiv|day)* and *H (day|indiv)* represent the conditional entropy given by

$$H(X|Y) = \sum_{y \in Y} p(y) \times H(X|Y = y)$$

$DOL_{day}$ measures daily specialization in task performance, i.e., how spread out the work is amongst individuals on a typical day while $DOL_{indiv}$ measures individual specialization across days, i.e., whether the same group of individuals contribute similarly to the task over days. Both range from 0 to 1, with 1 indicating that the task is performed only by a single individual on any given day in the case of $DOL_{day}$ and that the task is performed by a non-overlapping set of individuals on each day in the case of $DOL_{indiv}$. We thus defined our final metric of whether the task is consistently performed by the same individuals as

$$Individual\ consistency\ in\ task\ performance = 1 - DOL_{indiv}$$

For caregivers we obtained the identity of each caregiver, and the duration of care provided to the injured ant per day and combined data from five experimental days to calculate this metric. For foragers, we obtained the identity of each ant and the duration of time it spent in the foraging arena (obtained from the number of times it was detected in the foraging arena) during the experimental phase to calculate this metric. We obtained this consistency for each colony and task and then compared between tasks (M24).

*Caregivers who encounter injured ants*

Our observations revealed that individual caregivers show very low consistency over days, and most caregivers provided care only on one day. This would suggest that caregivers are not a specialized group. However, another possibility is that caregivers are specialized but individuals might not provide care on particular days due to a lack of opportunities to engage in care provisioning. To test whether this was the case, we first shortlisted all caregivers who encountered the injured ants in the experimental phase. We then obtained the proportion of these caregivers who provided care and compared this with the proportion of caregivers who did not provide care to that injured ant (M25).

*Caregiver characteristics*

We performed two types of comparisons to characterize caregivers: one focused on understanding their role within the social structure of the colony and another focused on understanding why they provide care on particular days and not others. In the first comparison, we asked where caregivers lie within the typical nurse/forager social structure[28,30] by comparing their properties with those of nurses and foragers. In the second comparison, to understand why caregivers provided care on particular days we compared their daily properties with those of two other groups: non-caregivers who encountered the injured ant but did not provide care and caregivers who did not provide care on that particular day but provided care on other days. We characterized these four groups as follows:

a. Nurses were identified based on their social maturity value[29,30]. Social maturity values range from 0 to 1, with values close to 0 indicating a strong association with the queen and hence the nurse community in the social network. We used interactions between all ant workers from the five baseline days before the manipulations to determine each ant's social maturity, and shortlisted individuals with a social maturity value of less than 0.2 as nurses.
b. Foragers were identified based on their presence in the foraging arena. We tweaked our previous definition of foragers (present at least on one day in the foraging arena) to be more conservative and only included individuals as foragers if they were present in the foraging arena during the 6-hour experimental phase on at least three out of five experimental days. Furthermore, if an individual identified as a forager at the colony level (it was present in the foraging arena on at least three out of five days) was not present in the foraging arena on a particular day, it was not considered as part of the forager group for that day.
c. Non-caregivers were identified based on their encounters with the injured ant in the experimental phase. For each caregiver ant, we calculated the encounters it had with the injured ant on each experimental day during the Experimental phase. To identify a group of non-caregivers, we then selected all other ants that, during the same time window, had a similar number of encounters with the injured ant. Non-caregivers were classified as such if their number of encounters ($E_n$) was within $E_{Ci} \pm E_{Colony}$ where $E_{Ci}$ is the number of encounters of the $i^{th}$ caregiver and $E_{Colony}$ is the average number of encounters of all ants in the colony with the injured ants across all five experimental phases. We used the colony-specific threshold of $E_{Colony}$ to account for differences between colonies in encounter rates. Thus, non-caregivers were nestmates who had similar opportunities to provide care as caregivers during a given Experimental phase but did not do so. One of the sterile injured ants was not detected by the tracking system in the Experimental phase, and we could not quantify

the encounters of nestmates with this injured ant. Some caregivers had a low number of encounters, and the range of value for $E_{Ci} \pm E_{Colony}$ could include 0. In this case we removed any non-caregivers with an encounter value of 0 as they did not meet the injured ant and would not have had any opportunity to provide care.

d. Caregivers who did not provide care on that particular day were the subset of all the caregivers identified in the colony over the five experimental days who did not provide care on that day.

We did not quantify caregivers and the other groups for injured ants which did not receive any care during the experimental phase (three sterile injured ants and three infected ants).

We compared movement related properties, social network profiles and parameters linked to the interactions with injured ants from the Pre-experimental phase between the caregivers and these groups. We focused on the Pre-experimental phase and not the Experimental phase to distinguish between characteristics that differentiate these groups under undisturbed conditions rather than characteristics that change as a result of care provision (e.g., by providing care, caregivers may interact more with only the injured ant in the Experimental phase, leading to a change in their activity or social network profile). The properties we compared are:

1. Average displacement (in pixels) per minute.
2. Proportion of time spent in the nest obtained by dividing the number of trajectory points in the nest arena by the total number of trajectory points of the ant.
3. Nest space use entropy which quantifies the evenness of the nest arena space use. To measure this, we divided the nest into 195 equally spaced hexagonal grids and calculated the number of times each ant was detected in each grid cell at 5Hz. We then obtained the proportion of detections in each grid cell and calculated the Shannon's entropy of this distribution (henceforth referred to as the nest space use entropy (50) for each ant in each phase as:

$$Nest\ space\ use\ entropy = -\sum_{i=1}^{195} (proportion_i * log(proportion_i))$$

   where i = 1 to 195 indicates each grid cell. Higher values of nest space use entropy indicate more even use of the nest arena whereas lower values indicate selective use of space. We removed ants which had non-zero values in 10 or fewer cells to filter out ants which were either mostly immobile or had very few detections.
4. Betweenness in the social network, a measure of centrality which quantifies the number of shortest paths between two pairs of nodes that pass through a given node.
5. Degree, which measures how many individuals each ant interacts with.
6. Strength, which quantifies the weighted sum of interactions for each ant. In our networks the connections were weighted based on the number of interactions between ants in a given time period.
7. Space use overlap quantified using an overlap index which measures how much the space use of individual nestmates overlaps with the space use of the injured ants based on their trajectories. To do so, we first obtained the mean X and Y coordinates per second for all caregivers, non-caregivers and injured ants. To ensure that we had sufficient trajectory data for estimating overlaps, we only selected data from arenas (nest or foraging) in which injured ants were present for at least ten minutes in each phase. Next, we selected caregivers and non-caregivers who were present in the

same arena as the injured ants for at least ten minutes and spent at least five minutes during each phase in these arenas. Then, we created a template raster layer using the injured individuals' trajectory data in each arena and obtained the kernel density estimate of an individual ant's space use based on this template raster. We then calculated the overlap at the 0.95 isopleth level between the space use of the injured ant and the caregivers and non-caregivers using a utilization distribution overlap index in each arena and averaged them across arenas for each individual.

8. Encounters with the injured ants in each phase.
9. Interactions with the injured ants in each phase.

We created one model per property with the ant type as the categorical response variable with five levels (M26 - M34).

*Prediction of care behavior*

Our characterization of caregivers indicated that the interaction between caregivers and injured ants is linked to care provision. To confirm this, we further evaluated whether an ant's behavior in the Pre-experimental phase could predict its propensity to provide care in the experimental phase. We calculated 13 individual behavioral properties including the nine described above and four additional properties in the Pre-experimental phase:

1. Total displacement (in pixels) over the whole phase.
2. Average number of interactions of each ant, which is the ratio between the strength and the degree of the ant.
3. The social maturity value of the ant, obtained from the five baseline days as described previously.
4. The network distance from the injured ants in the social network calculated as the number of edges in the shortest path connecting an ant with the injured ant.

In total we quantified four properties related to the movement (average and total displacement, proportion of time in the nest and nest space use entropy), five related to the social network profile (betweenness, degree, strength, average interactiveness and social maturity) and four related to the interactions with the injured ants (overlap index, encounters, interactions, and network distance) of all ants. For each type of property, we first scaled them based on the property type so that we could compare across experimental days, colonies, and properties. The experimental day was nested within colony for movement and network related properties. The injured ant type was nested within experimental day nested within colony for the injured ant related properties. Social maturity was an exception as we could only obtain one single value per ant and so we normalized this at the colony level. We then obtained the first two principal components of each property type - M1 and M2 for movement, N1 and N2 for network and A1 and A2 for injured ant related properties. We built 12 different models to determine which factors predicted an ant's probability to provide care in the experimental phase. The models included a combination of the different principal components and the z-normalized properties we calculated for each ant as predictors (M35 to M46). The models were compared using the Bayesian Information Criteria (BIC) to determine the model that best fit our data. We then used the predictors in the best fitting model (M42) to also determine if they predicted the amount of care ants provided in the Experimental phase (M47). For this analysis, we combined data from both experiments and included all ants from experimental days where injured individuals received care. In total, we had 136 individuals who provided care and 3277 that did not, which includes multiple values per ant corresponding to the number of experimental days per colony.

As an additional analysis, we performed the same comparison of the role of A1 and A2 in predicting care behavior (M48, M49), but only including data from caregivers and non-caregivers rather than from all ants. This allowed us to check whether A1 and A2 had the same predictive power even after controlling for the fact that only a subset of ants in the colony would have encountered the injured ant in the Experimental phase and hence had opportunities to provide care.

### Supplementary Results

Table S2: Structures of all the (generalized) linear models used in the statistical analyzes including the response variable, fixed and random effects, residual distribution, and the colonies from which the data is obtained. SI = sterile injured, II = infected injured.

| Model | Response variable | Fixed effects[1] | Random effects[2] | Error distribution | Colonies | Results |
|---|---|---|---|---|---|---|
| M1 | Number of leg licking events | Group (control/SI) and Leg (treated/non-treated) | Random slopes of group and leg within the combination of colony (3) and experimental day (5) | Poisson | 1, 2 and 3 | Table S3 and S4 |
| M2 | Log of the total leg licking duration | Group (control/SI)*Leg (treated/non-treated) | Random slopes of group and leg within the combination of colony (3) and experimental day (5) | Gaussian | 1, 2 and 3 | Table S3 and S4 |
| M3 | Number of leg licking events | Group (control/SI/II) and Leg (treated/non-treated) | Random slopes of group and leg within the combination of colony (3) and experimental day (5) | Poisson | 4, 5 and 6 | Table S3 and S4 |
| M4 | Log of the total leg licking duration | Group (control/SI/II) *Leg (treated/non-treated) | Random slopes of group and leg within the combination of colony (3) and experimental day (5) | Gaussian | 4, 5 and 6 | Table S3 and S4 |
| M5 | Number of allogrooming events | Group (control/SI) | Random slopes of group within the combination of colony (3) and experimental day (5) | Poisson | 1, 2 and 3 | Table S3 and S4 |
| M6 | Log of the total allogrooming duration | Group (control/SI) | Random slopes of group within the combination of colony (3) and experimental day (5) | Gaussian | 1, 2 and 3 | Table S3 and S4 |
| M7 | Number of allogrooming events | Group (control/SI/II) | Random slopes of group within the combination of colony (3) and experimental day (5) | Poisson | 4, 5 and 6 | Table S3 and S4 |
| M8 | Log of the total allogrooming duration | Group (control/SI/II) | Random slopes of group within the combination of colony (3) and experimental day (5) | Gaussian | 4, 5 and 6 | Table S3 and S4 |

| | | | | | | |
|---|---|---|---|---|---|---|
| M9 | Number of trophallaxis events | Group (control/SI) | Random slopes of group within the combination of colony (3) and experimental day (5) | Poisson | 1, 2 and 3 | Table S3 and S4 |
| M10 | Log of the total trophallaxis duration | Group (control/SI) | Random slopes of group within the combination of colony (3) and experimental day (5) | Gaussian | 1, 2 and 3 | Table S3 and S4 |
| M11 | Number of trophallaxis events | Group (control/SI/II) | Random slopes of group within the combination of colony (3) and experimental day (5) | Poisson | 4, 5 and 6 | Table S3 and S4 |
| M12 | Log of the total trophallaxis duration | Group (control/SI/II) | Random slopes of group within the combination of colony (3) and experimental day (5) | Gaussian | 4, 5 and 6 | Table S3 and S4 |
| M13 | Log of the average displacement per minute over 6 hours | Group (control/SI)*Phase (Baseline, Pre-experimental/Experimental/Post-experimental) | Random intercept of colony (3) | Gaussian | 1, 2 and 3 | Table S5 and S6 |
| M14 | Log of the average displacement per minute over 6 hours | Group (control/SI/II) *Phase (Baseline/Pre-experimental/Experimental/Post-experimental) | Random intercept of colony (3) | Gaussian | 4, 5 and 6 | Table S5 and S6 |
| M15 | Social maturity | Group (control/SI) | Random intercept of colony (3) | Beta[3] | 1, 2 and 3 | Section "Social Maturity of Focal Ants" |
| M16 | Social maturity | Group (control/SI/II) | Random intercept of colony (3) | Beta[3] | 4, 5 and 6 | Section "Social Maturity of Focal Ants" |
| M17 | Number of nodes | Network layer (wound care/allogrooming/trophallaxis) | Random intercept of colony (6) | Poisson | 1, 2, 3, 4, 5 and 6 | Section "Behavioral multiplex network |

| | | | | | | |
|---|---|---|---|---|---|---|
| | | | | | | comparison ” |
| M18 | Network layer overlap | Pairs of network layers (wound care-grooming/wound care-trophallaxis/grooming-trophallaxis) | Random intercept of colony (6) | Beta | 1, 2, 3, 4, 5 and 6 | Table S7 |
| M19 | Log median of the latency distribution | Behavior (wound care, grooming, trophallaxis) | Random intercept of colony (3) | Gaussian | 1, 2 and 3 | Table S8 and S9 |
| M20 | Log median of the latency distribution | Group (control/SI)*Behavior (allogrooming/trophallaxis) | Random intercept of colony (3) | Gaussian | 1, 2 and 3 | Table S8 and S9 |
| M21 | Log median of the latency distribution | Group (SI/II)*Behavior(wound care, allogrooming, trophallaxis) | Random intercept of colony (3) | Gaussian | 4, 5 and 6 | Table S8 and S9 |
| M22 | Log median of the latency distribution | Group (control/SI/II) *Behavior (allogrooming/trophallaxis) | Random intercept of colony (3) | Gaussian | 4, 5 and 6 | Table S8 and S9 |
| M23 | Log median of the latency distribution | Event type (woundcare/encounter) | Random intercept of replicate (2) nested within experimental day (5) nested within colony (6) | Gaussian | 1, 2, 3, 4, 5 and 6 | Section “Encounter based simulations of wound care” |
| M24 | Individual consistency in task performance | Task (foraging/wound care) | Random intercept of colony (6) | Beta | 1, 2, 3, 4, 5 and 6 | Main text |
| M25 | Proportion of caregivers | Caregiver type (encountered ant and provided care/encountered ant and did not provide care) | Random intercept of replicate (2) nested within experimental day (5) nested within colony (6) | Beta | 1, 2, 3, 4, 5 and 6 | Main text |

| | | | | | | |
|---|---|---|---|---|---|---|
| M26 | Log of the average displacement per minute over 6 hours in Pre-experimental phase | Ant type (caregiver/nurse/forager/non_caregiver/non-care provision caregiver) | Random intercept of experimental day (5) nested within colony (6) | Gaussian | 1, 2, 3, 4, 5 and 6 | Table S10 |
| M27 | Proportion of time spent in the nest in Pre-experimental phase | Ant type (caregiver/nurse/forager/non_caregiver/non-care provision caregiver) | Random intercept of experimental day (5) nested within colony (6) | Beta[3] | 1, 2, 3, 4, 5 and 6 | Table S10 |
| M28 | Space use entropy in the Pre-experimental phase | Ant type (caregiver/nurse/forager/non_caregiver/non-care provision caregiver) | Random intercept of experimental day (5) nested within colony (6) | Gaussian | 1, 2, 3, 4, 5 and 6 | Table S10 |
| M29 | Betweenness in the Pre-experimental phase | Ant type (caregiver/nurse/forager/non_caregiver/non-care provision caregiver) | Random intercept of experimental day (5) nested within colony (6) | Gamma[4] | 1, 2, 3, 4, 5 and 6 | Table S10 |
| M30 | Degree in the Pre-experimental phase | Ant type (caregiver/nurse/forager/non_caregiver/non-care provision caregiver) | Random intercept of experimental day (5) nested within colony (6) | Negative binomial | 1, 2, 3, 4, 5 and 6 | Table S10 |
| M31 | Strength in the Pre-experimental phase | Ant type (caregiver/nurse/forager/non_caregiver/non-care provision caregiver) | Random intercept of experimental day (5) nested within colony (6) | Negative binomial | 1, 2, 3, 4, 5 and 6 | Table S10 |
| M32 | Space use overlap with injured ants in the Pre-experimental phase | Ant type (caregiver/nurse/forager/non_caregiver/non-care provision caregiver) | Random intercept of experimental day (5) nested within colony (6) | Gaussian | 1, 2, 3, 4, 5 and 6 | Table S10 |
| M33 | Encounters with the injured ants in the Pre- | Ant type (caregiver/nurse/forager/non_caregiver/no | Random intercept of replicate (2) nested within | Negative binomial | 1, 2, 3, 4, 5 and 6 | Table S10 |

| | | | | | | |
|---|---|---|---|---|---|---|
| | experimental phase | n-care provision caregiver) | experimental day (5) nested within colony (6) | | | |
| M34 | Interactions with injured ants in the Pre-experimental phase | Ant type (caregiver/nurse/forager/non_caregiver/non-care provision caregiver) | Random intercept of replicate (2) nested within experimental day (5) nested within colony (6) | Negative binomial | 1, 2, 3, 4, 5 and 6 | Table S10 |
| M35 | Care provision in Experimental phase (binary variable) | Z-normalized individual properties (average displacement + total displacement + space use entropy + proportion of time in nest + betweenness + degree + strength + average number of interactions + overlap index + encounters + interactions + network distance from the injured ant) in the Pre-experimental phase | Random intercept of experimental day (5) nested within colony (6) | Binomial | 1, 2, 3, 4, 5 and 6 | Table S12 |
| M36 | Care provision in Experimental phase (binary variable) | Principal Components of all property categories (M1 + M2 + N1 + N2 + A1 + A2) in the Pre-experimental phase | Random intercept of experimental day (5) nested within colony (6) | Binomial | 1, 2, 3, 4, 5 and 6 | Table S12 |
| M37 | Care provision in Experimental phase (binary variable) | Z-normalized movement related properties (average displacement + total displacement + space use entropy) in the Pre-experimental phase | Random intercept of experimental day (5) nested within colony (6) | Binomial | 1, 2, 3, 4, 5 and 6 | Table S12 |
| M38 | Care provision in Experimental | Z-normalized network related properties (betweenness + degree + strength + | Random intercept of experimental day (5) nested within colony (6) | Binomial | 1, 2, 3, 4, 5 and 6 | Table S12 |

| | | | | | | |
|---|---|---|---|---|---|---|
| | phase (binary variable) | average number of interactions) in the Pre-experimental phase | | | | |
| M39 | Care provision in Experimental phase (binary variable) | Z-normalized injured ant affinity related properties (overlap index + encounters + interactions + network distance from the injured ant) in the Pre-experimental phase | Random intercept of experimental day (5) nested within colony (6) | Binomial | 1, 2, 3, 4, 5 and 6 | Table S12 |
| M40 | Care provision in Experimental phase (binary variable) | Principal components of movement related properties (M1 + M2) in the Pre-experimental phase | Random intercept of experimental day (5) nested within colony (6) | Binomial | 1, 2, 3, 4, 5 and 6 | Table S12 |
| M41 | Care provision in Experimental phase (binary variable) | Principal components of network related properties (N1 + N2) in the Pre-experimental phase | Random intercept of experimental day (5) nested within colony (6) | Binomial | 1, 2, 3, 4, 5 and 6 | Table S12 |
| M42 | Care provision in Experimental phase (binary variable) | Principal components of injured ant affinity related properties (A1 + A2) in the Pre-experimental phase | Random intercept of experimental day (5) nested within colony (6) | Binomial | 1, 2, 3, 4, 5 and 6 | Table S12 and S13 |
| M43 | Care provision in Experimental phase (binary variable) | Z-normalized interactions with the injured ants in the Pre-experimental phase | Random intercept of experimental day (5) nested within colony (6) | Binomial | 1, 2, 3, 4, 5 and 6 | Table S12 |
| M44 | Care provision in Experimental phase (binary variable) | Z-normalized encounters with the injured ants in the Pre-experimental phase | Random intercept of experimental day (5) nested within colony (6) | Binomial | 1, 2, 3, 4, 5 and 6 | Table S12 |

| | | | | | | |
|---|---|---|---|---|---|---|
| M45 | Care provision in Experimental phase (binary variable) | Z-normalized overlap index in the Pre-experimental phase | Random intercept of experimental day (5) nested within colony (6) | Binomial | 1, 2, 3, 4, 5 and 6 | Table S12 |
| M46 | Care provision in Experimental phase (binary variable) | Z-normalized social maturity | Random intercept of experimental day (5) nested within colony (6) | Binomial | 1, 2, 3, 4, 5 and 6 | Table S12 |
| M47 | Total Number of wound care events in Experimental phase | Principal components of injured ant affinity related properties (A1 + A2) in the Pre-experimental phase | Random intercept of experimental day (5) nested within colony (6) | Negative binomial | 1, 2, 3, 4, 5 and 6 | Table S13 |
| M48 | Care provision in Experimental phase (binary variable) | Principal components of injured ant affinity related properties (A1 + A2) in the Pre-experimental phase | Random intercept of experimental day (5) nested within colony (6) | Binomial | 1, 2, 3, 4, 5 and 6 (only including caregivers and non-caregivers) | Table S13 |
| M49 | Total Number of wound care events in Experimental phase | Principal components of injured ant affinity related properties (A1 + A2) in the Pre-experimental phase | Random intercept of experimental day (5) nested within colony (6) | Negative binomial | 1, 2, 3, 4, 5 and 6 (only including caregivers and non-caregivers) | Table S13 |

1. For fixed effects which are categorical, the levels of the variable are provided in brackets. An interaction between the two fixed effects terms is denoted by an '*'.
2. For random effects, the numbers in brackets represent the number of levels of the random effect variable. In most models, we started with a maximal random effect structure but had to fit a reduced one due to convergence issues (e.g., in model M13 we could not fit the random intercept of experimental day nested within colony).
3. For beta residual distributions, the response variable has to lie in (0, 1). Since proportion and social maturity values lie in [0, 1], we added a small value to modify the original values for running the model.
4. To fit the gamma error structure, we had to remove individuals which had a betweenness value of zero (69 out of 2261 individual values).

### Survival analyses

#### Test of *Pseudomonas aeruginosa* concentration lethality

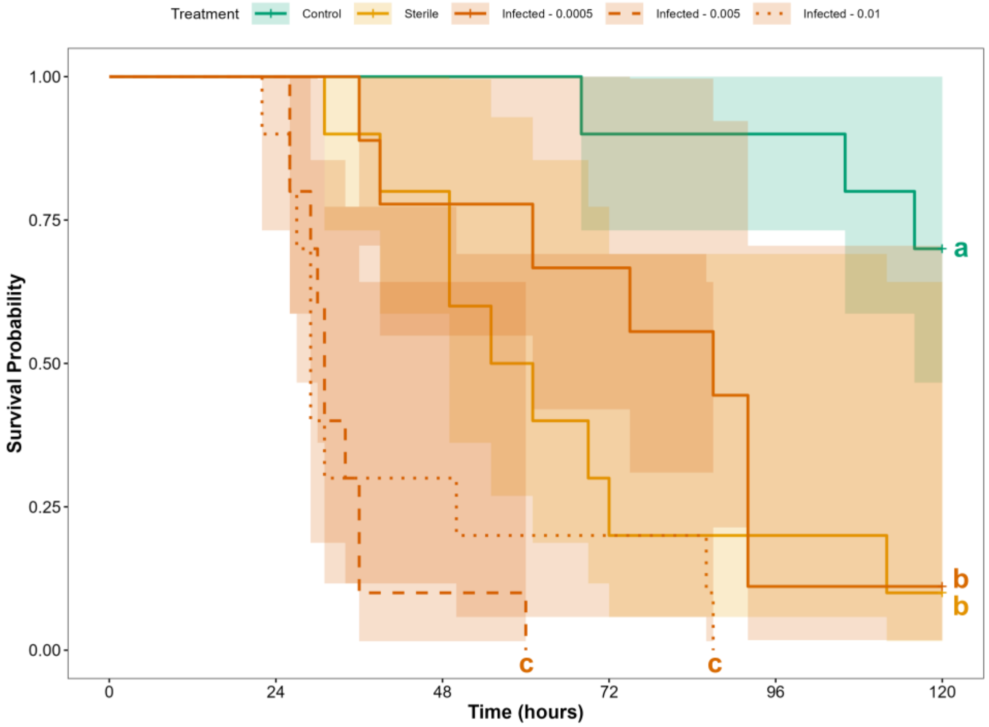


Figure S1: Survival probabilities and their confidence limits of ants (n = 10 per group) exposed to three different concentrations (optical densities) of *P. aeruginosa* (Infected - 0.0005, Infected - 0.005, Infected - 0.01), ants treated with a sterile wound (Sterile) and uninjured ants (Control) over time. Colors of the lines denote injury and infection status with green for uninjured control ants, orange for sterile injured ants and red for infected injured ants and the line type denotes the concentration for the infected ants (solid = 0.0005, dashed = 0.005, dotted = 0.01). Concentration OD = 0.005 was used in the subsequent experiments. Different letters at the end of the lines denote whether the focal groups were significantly different from each other. Plus symbols at the end of the lines indicate that individuals were still alive after 120 hours of observation.

We found that ants infected with the smallest bacterial concentration (0.0005 OD) did not differ in survival probability from uninfected injured ants (cox model coefficient = 0.28, CI = 0.50-3.47, p = 0.57), but had shorter survival times than uninjured Control ants (cox model coefficient = -1.46, CI = 0.07 - 0.80, p = 0.02), while having higher survival than ants infected with higher bacterial concentrations (0.0005 vs 0.005: cox model coefficient = 2.01, CI = 2.55 - 21.73, p < 0.001; 0.0005 vs 0.01: cox model coefficient = 1.50, CI = 1.68 - 11.84, p = 0.003). Ants infected with a 0.005 OD solution had significantly shorter survival times than Control (cox model coefficient = -3.46, CI = 0.008 - 0.121, p < 0.001) and sterile injured ants (cox model coefficient = -1.73, CI = 0.006 - 0.494, p < 0.001). These latter results are in line with previous work using the same bacteria but in *C. floridanus* ants [13]. The survival of ants with injuries infected with a 0.005 OD solution did not differ from ants infected with a 0.01 OD solution (cox model coefficient = -0.51, CI = 0.232 - 0.1543, p = 0.29, Fig S1). Given these results, we decided to use a 0.005 dilution.

#### Infection survival experiment

All three groups of ants tested in isolation (Control, sterile injured and infected injured) differed from each other in their survival, with Control ants having the highest survival probability at any given point in time (Control vs sterile injured: cox model coefficient = 1.31, CI = 1.58 - 8.69, p = 0.003; Control vs infected injured: cox model coefficient = 2.08, CI = 3.28-19.56, p < 0.001). Sterile injured ants had overall higher survival probability than infected injured ants (sterile vs infected: cox model coefficient = 0.77, CI = 1.03-4.51, p = 0.04, Fig S2) indicating that the infections indeed increased mortality when infected ants did not receive care.

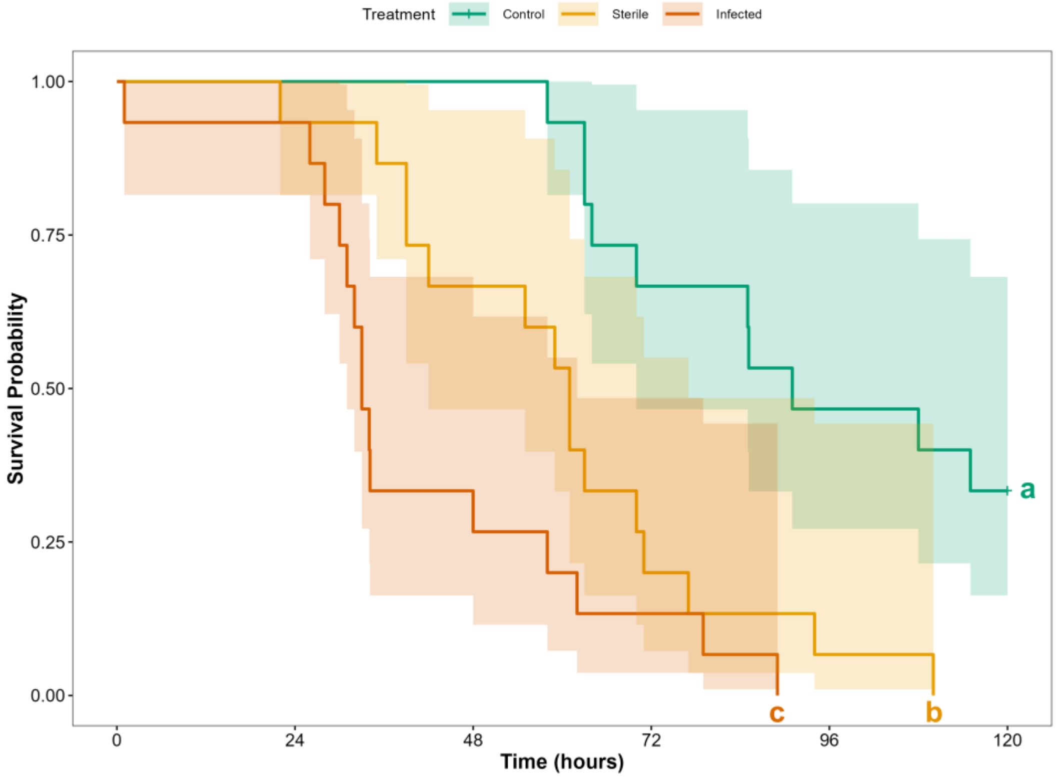


Figure S2: Survival probabilities and their confidence limits of isolated ants injured with infected wounds of 0.005 OD dilutions of *P. aeruginosa* (Infected, red), ants treated with a sterile wound (Sterile, orange) and uninjured ants (Control, green) over time. Different letters at the end of the lines denote whether the focal groups (n = 15 per group) were significantly different from each other. Plus symbols at the end of the lines indicate that individuals were still alive after 120 hours of observation.

### Care provisioning and food provisioning

#### Behavioral response to focal ants

In colonies 1 to 3, where only sterile injured and control ants were present, we found independent effects of the leg (i.e., treated leg or non-treated legs) and focal group on the duration of leg licking. Specifically, licking events were longer when directed towards injured ants compared to the control ants regardless of which type of leg was licked (estimate ± SE = -3.17 ± 0.73, p < 0.001) and when treated legs were licked (i.e., wound care in case of sterile and infected injured ants) compared to when licking non-treated legs (estimate= -1.30 ± 0.58, p = 0.03). This is in slight contrast to the frequency results, where we found a significant interaction between which leg was licked and focal group (see main text). Similarly, in colonies 4 to 6, both sterile and infected ants received longer wound care bouts than the licking bouts control ants received on the equivalent treated leg (chi-squared = 59.543, df = 5, $p < 0.001$). Non-treated legs of infected ants were also licked longer than the non-treated legs of control ants, which was not the case for ants with sterile injuries. Ants with sterile and infected injuries did not differ in the duration of cleaning of both treated and non-treated legs (Table S3 and S4).

In all colonies, control ants received significantly fewer allogrooming events than injured ants (Table S4). However, ants with sterile and infected wounds did not differ (Table S4). In terms of behavioral duration, infected and sterile injured received longer allogrooming events than control ants (Table S4). Sterile ants did not differ from control ants or infected ants in terms of allogrooming duration (Table S4). The three focal groups never differed in their trophallaxis frequency or duration.

Table S3: Estimated marginal means and confidence intervals (CI) of the total count and duration of each behavior for each type of focal ant in the Experimental phase. Wound care is equivalent to licking the treated (injured) leg in injured ants.

| Data type | Colonies | Behavior | Control | | SI | | II | |
|---|---|---|---|---|---|---|---|---|
| | | | Estimate | CI | Estimate | CI | Estimate | CI |
| Count | 1, 2 and 3 | Wound care | 1.25 | 0.65 - 2.42 | 20.94 | 12.15 - 36.09 | | |
| | | Licking non-treated legs | 1.95 | 1.04 - 3.66 | 4.33 | 2.42 - 7.77 | | |
| | | Allogrooming | 6.99 | 4.26 - 11.46 | 22.17 | 13.7 - 35.86 | | |
| | | Trophallaxis | 14.55 | 10.50 - 20.16 | 22.86 | 16.27 - 32.11 | | |
| | 4, 5 and 6 | Wound care | 0.03 | 0.003 - 0.22 | 10.90 | 4.65 - 25.59 | 6.92 | 3.74 - 12.83 |
| | | Licking non-treated legs | 0.41 | 0.18 - 0.96 | 1.31 | 0.52 - 3.27 | 2.93 | 1.54 - 5.58 |
| | | Allogrooming | 1.59 | 0.91 - 2.75 | 9.06 | 4.87 - 16.85 | 8.45 | 5.28 - 13.54 |
| | | Trophallaxis | 6.81 | 3.74 - 12.38 | 4.51 | 2.21 - 9.20 | 7.90 | 4.35 - 14.34 |

| | | | | | | | | |
|---|---|---|---|---|---|---|---|---|
| Duration | 1, 2 and 3 | Wound care | 2.54 | 1.50 - 3.57 | 5.71 | 4.64 - 6.79 | | |
| | | Licking non-treated legs | 2.55 | 1.51 - 3.58 | 4.41 | 3.33 - 5.48 | | |
| | | Allogrooming | 4.49 | 3.45 - 5.54 | 5.86 | 4.77 - 6.94 | | |
| | | Trophallaxis | 5.95 | 5.46 - 6.43 | 6.44 | 5.93 - 6.94 | | |
| | 4, 5 and 6 | Wound care | 1.5e-15 | -0.95 - 0.95 | 5.21 | 4.10 - 6.32 | 4.50 | 3.55 - 5.45 |
| | | Licking non-treated legs | 1.45 | 0.50 - 2.40 | 3.65 | 2.55 - 4.76 | 3.67 | 2.72 - 4.62 |
| | | Allogrooming | 2.97 | 2.06 - 3.89 | 5.51 | 4.59 - 6.43 | 5.26 | 4.35 - 6.18 |
| | | Trophallaxis | 4.81 | 3.68 - 5.93 | 4.85 | 3.72 - 5.98 | 4.83 | 3.71 - 5.96 |

Table S4: Contrasts between focal groups and between specific behaviors of the total count and duration of each behavior during the Experimental phase. SI = sterile injured, II = infected injured.

| Data type | Colonies | Comparison | Level | Difference | t/z value | p value |
|---|---|---|---|---|---|---|
| Count | 1, 2 and 3 | Control vs SI | Wound care | -2.83 | -6.03 | < 0.001 |
| | | | Licking other, non-injured legs | -0.81 | -1.72 | 0.31 |
| | | | Allogrooming | -1.15 | -3.41 | < 0.001 |
| | | | Trophallaxis | -0.45 | -1.70 | 0.089 |
| | | Wound care vs licking non-treated legs | Control | -0.45 | -1.72 | 0.31 |
| | | | SI | 1.57 | 8.97 | < 0.001 |
| | 4, 5 and 6 | Control vs SI | Wound care | -5.97 | -5.23 | < 0.001 |

| | | | | | | |
|---|---|---|---|---|---|---|
| | | | Licking other, non-injured legs | -1.15 | -1.77 | 0.48 |
| | | | Allogrooming | -1.74 | -4.22 | < 0.001 |
| | | | Trophallaxis | 0.41 | 1.19 | 0.457 |
| | | Control vs II | Wound care | -5.51 | -5.50 | < 0.001 |
| | | | Licking other, non-injured legs | -1.96 | -6.08 | < 0.001 |
| | | | Allogrooming | -1.67 | -9.59 | < 0.001 |
| | | | Trophallaxis | -0.15 | -1.47 | 0.306 |
| | | SI vs II | Wound care | 0.45 | 0.82 | 0.960 |
| | | | Licking other, non-injured legs | -0.81 | -1.37 | 0.740 |
| | | | Allogrooming | 0.07 | 0.19 | 0.981 |
| | | | Trophallaxis | -0.56 | -1.63 | 0.232 |
| | | Wound care vs licking non-treated legs | Control | -2.70 | -2.51 | 0.12 |
| | | | SI | 2.12 | 7.05 | < 0.001 |
| | | | II | 0.86 | 2.99 | 0.030 |
| Duration | 1, 2 and 3 | Control vs SI | Cleaning legs* | -3.17 | | < 0.001 |
| | | | Allogrooming | 183 | 0.88 | 0.390 |
| | | | Trophallaxis | -0.49 | -1.46 | 0.163 |

| | | | | | | |
|---|---|---|---|---|---|---|
| | | Wound care vs licking non-treated legs* | | 1.30 | | 0.030 |
| | 4, 5 and 6 | Control vs SI | Wound care | -5.21 | -6.63 | < 0.001 |
| | | | Licking non-treated legs | -2.20 | -2.80 | 0.080 |
| | | | Allogrooming | -2.54 | -4.42 | < 0.001 |
| | | | Trophallaxis | -0.04 | -0.06 | > 0.998 |
| | | Control vs II | Wound care | -4.50 | -7.19 | < 0.001 |
| | | | Licking non-treated legs | -2.22 | -3.55 | 0.010 |
| | | | Allogrooming | -2.29 | -3.99 | 0.003 |
| | | | Trophallaxis | -0.02 | -0.04 | > 0.999 |
| | | SI vs II | Wound care | 0.71 | 0.90 | 0.940 |
| | | | Licking non-treated legs | -0.02 | -0.02 | > 0.999 |
| | | | Allogrooming | 0.24 | 0.43 | 0.910 |
| | | | Trophallaxis | 0.02 | 0.02 | > 0.999 |
| | | Wound care vs licking non-treated legs | Control | -1.45 | -2.28 | 0.220 |
| | | | SI | 1.55 | 2.43 | 0.160 |
| | | | II | 0.82 | 1.29 | 0.790 |

* In the behavioral duration data for colonies 1 to 3, there was no significant interaction effect between the type of leg (injured leg vs non-injured legs) and focal group (control vs sterile ants), so we provide the contrasts from the main effects only.

#### Activity of focal individuals

Focal ants did not differ in their activity levels in the Baseline, Pre-experimental, Experimental or Post-experimental phases (Table S5 and S6). Thus, differences in the behaviors received by the focal ants were not due to sterile and infected injured ants being more active than control ants.

Table S5: Estimated marginal means and confidence intervals (CI) of average displacement per minute (pixels per minute) for each focal group.

| Colonies | Phase | Control | | Sterile injured | | Infected injured | |
|---|---|---|---|---|---|---|---|
| | | Estimate | CI | Estimate | CI | Estimate | CI |
| 1, 2 and 3 | Baseline | 863.81 | 515.77 - 1446.71 | 778.38 | 464.76 - 1303.62 | | |
| | Pre-experimental | 746.17 | 445.53 - 1249.68 | 874.30 | 517.22 - 1477.88 | | |
| | Experimental | 773.74 | 461.99 - 1295.85 | 873.04 | 521.28 - 1462.16 | | |
| | Post-experimental | 662.46 | 395.55 - 1109.49 | 604.11 | 349.22 - 1045.04 | | |
| 4, 5 and 6 | Baseline | 774.59 | 469.97 - 1276.64 | 542.13 | 328.93 - 893.51 | 918.40 | 557.23 - 1513.66 |
| | Pre-experimental | 543.14 | 329.54 - 895.17 | 558.71 | 338.99 - 920.83 | 797.44 | 483.84 - 1314.30 |
| | Experimental | 657.67 | 399.04 - 1083.94 | 583.40 | 348.36 - 976.99 | 858.79 | 521.06 - 1415.41 |
| | Post-experimental | 445.77 | 270.47 - 734.69 | 592.02 | 314.91 - 1112.99 | 433.97 | 244.07 - 771.62 |

Table S6: Contrasts between focal groups of their average displacement per minute. SI = sterile injured, II = infected injured.

| Colonies | Focal groups | Phase | Difference | t value | p value |
|---|---|---|---|---|---|
| 1, 2 and 3 | Control vs SI | Baseline | 0.10 | 0.38 | 0.707 |
| | | Pre-experimental | -0.16 | -0.56 | 0.574 |
| | | Experimental | -0.12 | -0.44 | 0.663 |

| | | | | | |
|---|---|---|---|---|---|
| | | Post-experimental | 0.09 | 0.31 | 0.754 |
| 4, 5 and 6 | Control vs SI | Baseline | 0.36 | 1.05 | 0.590 |
| | | Pre-experimental | -0.03 | 0.08 | 0.934 |
| | | Experimental | 0.12 | 0.35 | 0.866 |
| | | Post-experimental | -0.28 | -0.72 | > 0.999 |
| | Control vs II | Baseline | -0.17 | 0.50 | 0.617 |
| | | Pre-experimental | -0.38 | -1.13 | 0.779 |
| | | Experimental | -0.27 | -0.79 | 0.866 |
| | | Post-experimental | 0.03 | 0.07 | > 0.999 |
| | SI vs II | Baseline | 0.53 | 1.55 | 0.367 |
| | | Pre-experimental | 0.36 | 1.05 | 0.779 |
| | | Experimental | 0.39 | 1.12 | 0.795 |
| | | Post-experimental | -0.31 | -0.74 | > 0.999 |

#### Social maturity of focal ants

Control and sterile injured ants did not differ significantly in terms of social maturity (estimated marginal means and confidence intervals of social maturity of control ants = 0.88 [0.71 - 0.95], sterile injured ants = 0.91 [0.77 - 0.97]; difference estimate for control vs sterile injured = -0.36, $z = -1.21$, $p = 0.225$). Sterile injured and infected ants did not differ from control ants (estimated marginal means and confidence intervals of social maturity of control ants = 0.72 [0.13 - 0.98], sterile injured ants ants = 0.73 [0.14 - 0.98], infected ants = 0.70 [0.12 - 0.98]; difference estimate for control vs sterile injured ants = -0.07, $z = -0.22$, $p > 0.999$, control vs infected = 0.09, $z = 0.28$, $p > 0.999$, sterile injured ants vs infected = 0.15, $z = 0.50$, $p > 0.999$). Thus, differences in the behavioral response of nestmates to the focal ants was not due to differences in the social maturity of the focal ants.

#### Behavioral multiplex network comparison

We found that wound care networks were significantly smaller than trophallaxis networks with allogrooming networks having an intermediate size (size of wound care layer = 20 ± 4.69, allogrooming = 35 ± 6, trophallaxis = 55.7 ± 14.7; wound care vs trophallaxis: difference = -1.02, $z = 9.62$, $p < 0.001$;

allogrooming vs trophallaxis : difference = -0.46, z = 5.27, p < 0.001; wound care vs allogrooming: difference = -40.56, z = -4.89, p < 0.001). Network layer overlap between the wound care and allogrooming layers was significantly higher than their overlap with the trophallaxis layer. Additionally, the network layer overlap between allogrooming-trophallaxis layers was higher than the network layer overlap between wound care-trophallaxis (Table S7). These results show that wound care and allogrooming are performed by similar individuals at the colony level in contrast with trophallaxis.

Table S7: Estimated marginal mean network layer overlap and confidence intervals (CI) of networks of different behaviors and contrasts between network pairs.

| Behavioral Layers/Comparison | Estimate | CI | Difference | z value | p value |
|---|---|---|---|---|---|
| Wound care-allogrooming | 0.39 | 0.32 – 0.47 | | | |
| Wound care-trophallaxis | 0.08 | 0.06 – 0.12 | | | |
| Allogrooming-trophallaxis | 0.14 | 0.10 – 0.19 | | | |
| Wound care-allogrooming vs wound care-trophallaxis | | | 1.94 | 8.59 | 0.0001 |
| Wound care-allogrooming vs allogrooming-trophallaxis | | | 1.41 | 7.10 | 0.0001 |
| Wound care-trophallaxis vs allogrooming-trophallaxis | | | -0.53 | -2.12 | 0.034 |

#### Correlations between behaviors at the individual level

When comparing behaviors at the individual level, the total duration of wound care and allogrooming performed by a nestmate towards the injured ant showed a significant positive correlation (Fig. S3A), while the total duration of wound care and trophallaxis did not (Pearson correlation coefficients and associated p values provided in Fig S3). Similarly, there was no significant correlation between the total duration of allogrooming and trophallaxis (Fig. S3C). These results show that behavioral separation between care and food provision we observed at the colony level is also reflected in individual behavior.

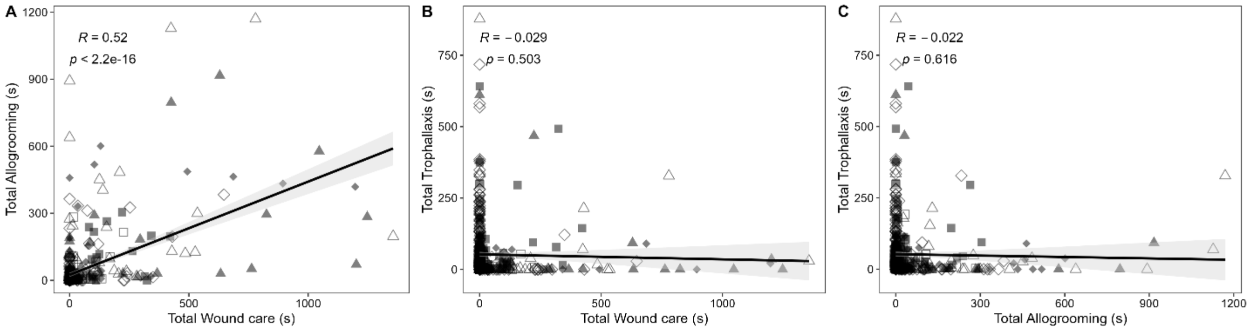


Figure S3: The correlations between the three different behaviors (wound care, allogrooming and trophallaxis) at the individual level. Each facet corresponds to the correlation between one pair of behaviors and the Pearson's correlation coefficient (R) and associated p value are provided in the text inset within the plot. Each point represents data from one colony with filled shapes representing data from colonies 1 to 3 and unfilled shapes representing data from colonies 4 to 6. The black line represents the regression line between the two variables with the shaded region the 95% confidence interval.

#### Behavioral dynamics over time

Comparing medians of the distribution of when the behavioral events occurred, we found that wound care and allogrooming were temporally linked (Table S8 and S9). In colonies 1 to 3, within sterile injured ants, the median time for wound care and allogrooming did not differ from each other and was significantly earlier than the median time for trophallaxis. Comparing between control and sterile ants, control ants had a significantly earlier median in allogrooming but did not differ from sterile ants in the median time for trophallaxis. In colonies 4 to 6, sterile and infected ants did not significantly differ from each other in the median time of any of the three behaviors. In both groups, wound care and allogrooming had similar medians and this was significantly earlier than the median time of trophallaxis (with the exception that in the sterile ants the median for wound care was earlier than the median for trophallaxis, but not significantly so with a $p = 0.075$). When compared to control ants, sterile and infected ants did not differ in the median time of allogrooming or trophallaxis.

Table S8: Average medians and confident intervals (CI) of the distributions of occurrence time of behavioral events in minutes per behavior and focal group in each experiment.

| Colonies | Behavior | Control | | Sterile injured | | Infected injured | |
|---|---|---|---|---|---|---|---|
| | | Estimate | CI | Estimate | CI | Estimate | CI |
| 1, 2 and 3 | Wound care | | | 67.67 | 45.89 – 99.78 | | |
| | Allogrooming | 40.34 | 28.44 – 57.23 | 78.62 | 54.09 – 114.27 | | |
| | Trophallaxis | 141.77 | 103.63 – 193.94 | 191.73 | 131.00 – 280.63 | | |
| 4, 5 and 6 | Wound care | | | 83.56 | 55.35 - 126.16 | 62.31 | 39.55 - 98.19 |
| | Allogrooming | 113.81 | 63.77 - 203.12 | 75.17 | 52.29 - 108.06 | 62.61 | 43.15 - 90.83 |
| | Trophallaxis | 130.72 | 84.84 - 201.41 | 150.21 | 96.05 - 234.93 | 180.87 | 117.83 - 277.64 |

Table S9: Contrasts between the time dynamics of the different behaviors within and across focal groups. Estimates represent differences of log (median) of the distribution of occurrence time of behavioral events. SI = sterile injured, II = infected injured

| Colonies | Focal groups | Behavior | Difference | t value/ratio | p value |
|---|---|---|---|---|---|
| 1, 2 and 3 | SI | Wound care vs allogrooming | -0.15 | -0.71 | 0.482 |
| | | Wound care vs trophallaxis | -1.04 | -4.85 | <0.001 |
| | | Allogrooming vs trophallaxis | -0.89 | -4.29 | 0.0002 |
| | Control vs SI | Allogrooming | -0.67 | -3.01 | 0.004 |
| | | Trophallaxis | -0.31 | -1.47 | 0.146 |
| 4, 5 and 6 | SI | Wound care vs allogrooming | 0.11 | 0.71 | 0.662 |
| | | Wound care vs trophallaxis | -0.59 | -2.12 | 0.075 |
| | | Allogrooming vs trophallaxis | -0.69 | -2.70 | 0.027 |
| | II | Wound care vs allogrooming | -0.004 | -0.02 | 0.986 |
| | | Wound care vs trophallaxis | -1.07 | -3.77 | 0.001 |
| | | Allogrooming vs trophallaxis | -1.06 | -4.22 | 0.0002 |
| | Control vs SI | Allogrooming | 0.41 | 1.34 | 0.368 |
| | | Trophallaxis | -0.14 | -0.51 | 0.929 |
| | Control vs II | Allogrooming | 0.60 | 1.92 | 0.178 |
| | | Trophallaxis | -0.33 | -1.24 | 0.656 |
| | SI vs II | Wound care | 0.29 | 1.04 | 0.301 |
| | | Allogrooming | 0.18 | 0.82 | 0.414 |
| | | Trophallaxis | -0.19 | -0.66 | 0.514 |

### Caregiver analyzes

#### Encounter based simulations of wound care

Our comparison between the median latencies of wound care events and encounters revealed that the median time of occurrence of wound care events was significantly earlier compared to the median time of occurrence of encounters (median time of occurrence of wound care events = 85.80 ± 54.70, encounters = 148.74 ± 65.25; difference = -62.94, $t = -4.19$, $p < 0.001$).

In the comparison between $S_{low}$, $S_{med}$, $S_{high}$ and $S_{empirical}$, we found that scenario $S_{low}$ came closest to the observed temporal dynamics of wound care, but all four scenarios overestimated the number of caregivers compared to our empirical observations (Fig. S4). Out of 30 ants, the empirical median time of the distribution of wound care events was significantly different from the expected values based on the simulations in 14 ants for scenario $S_{low}$, 22 ants for scenario $S_{med}$, 28 ants for scenario $S_{high}$ and 16 ants for $S_{empirical}$. In the case of number of caregivers, the empirical number of caregivers observed was significantly different from the expected values based on the simulations for all 30 ants in all scenarios.

Table S10: The median time (in minutes) of the distribution of wound care events from the Experimental phase and the number of caregivers obtained from our empirical observations and the simulations. For the data from the simulations, the values represent the median obtained from 1000 simulations for each ant and scenario. The values in the brackets represent the likelihood of the empirical value based on the null distribution of values obtained from the simulations.

| Injured Ant No | Empirical | | $S_{empirical}$ | | $S_{low}$ | | $S_{med}$ | | $S_{high}$ | |
|---|---|---|---|---|---|---|---|---|---|---|
| | Median Time | No. of caregivers | Median Time | No. of caregivers | Median Time | No. of caregivers | Median Time | No. of caregivers | Median Time | No. of caregivers |
| 1 | 45.84 | 4 | 99.13 (0.06) | 13 (0) | 43.33 (0.868) | 12.5 (0) | 6.74 (0) | 13 (0) | 4.2 (0) | 13 (0) |
| 2 | 70.87 | 5 | 120.81 (0.104) | 13 (0) | 37.72 (0.004) | 14 (0) | 5.29 (0) | 14.5 (0) | 3.43 (0) | 14.5 (0) |
| 3 | 82.06 | 5 | 172.51 (0.01) | 14.5 (0) | 99.08 (0.478) | 15.5 (0) | 22.18 (0) | 16 (0) | 12.8 (0) | 16 (0) |
| 4 | 10.89 | 3 | 12.58 (0.972) | 6.5 (0) | 13.33 (0.704) | 6.5 (0) | 9.45 (0.172) | 6.5 (0) | 7.37 (0) | 6.5 (0) |
| 5 | 165.54 | 3 | 81.36 (0.048) | 23.5 (0) | 85.35 (0.062) | 22.5 (0) | 22.42 (0) | 18 (0) | 11.8 (0) | 18.5 (0) |
| 6 | 132.92 | 3 | 20.9 (0) | 10.5 (0) | 24.12 (0) | 10.5 (0) | 5.22 (0) | 11 (0) | 3.91 (0) | 9.5 (0) |
| 7 | 34.74 | 4 | 163.84 (0) | 18.5 (0) | 153.14 (0) | 18 (0) | 47.8 (0.094) | 15 (0) | 17.99 (0.008) | 12 (0) |
| 8 | 70.54 | 5 | 167.41 (0) | 26.5 (0) | 172.08 (0) | 26 (0) | 167.82 (0) | 26 (0) | 165.06 (0) | 25.5 (0) |
| 9 | 147.97 | 3 | 272.9 (0.134) | 5.5 (0) | 91.18 (0.066) | 4.5 (0.018) | 13.83 (0) | 5 (0.01) | 6.91 (0) | 5.5 (0) |
| 10 | 28.2 | 2 | 231.93 (0) | 17.5 (0) | 200.61 (0) | 17.5 (0) | 27.67 (0.852) | 16.5 (0) | 9.97 (0) | 16 (0) |

| | | | | | | | | | |
|---|---|---|---|---|---|---|---|---|---|
| 11 | 15.34 | 2 | 257.89 (0.006) | 6.5 (0) | 255.9 (0.008) | 7 (0) | 14.78 (0.738) | 7.5 (0) | 5.3 (0.008) | 7.5 (0) |
| 12 | 75.94 | 9 | 159.74 (0.014) | 23.5 (0) | 100.19 (0.494) | 22.5 (0) | 16.96 (0) | 23 (0) | 9.18 (0) | 23 (0) |
| 13 | 81.32 | 10 | 117.64 (0.216) | 21 (0) | 117.56 (0.194) | 21 (0) | 31.06 (0) | 20 (0) | 10.35 (0) | 17.5 (0) |
| 14 | 77.52 | 4 | 151.02 (0.656) | 14 (0) | 69.55 (0.594) | 13 (0) | 18.21 (0) | 13 (0) | 12.97 (0) | 14 (0) |
| 15 | 302.46 | 7 | 131.35 (0) | 20 (0) | 132.57 (0) | 19.5 (0) | 14.39 (0) | 19.5 (0) | 6.03 (0) | 19.5 (0) |
| 16 | 73.47 | 10 | 170.93 (0) | 24 (0) | 162.29 (0) | 24 (0) | 83.15 (0.346) | 23.5 (0) | 61.9 (0) | 23 (0) |
| 17 | 51.27 | 4 | 210.6 (0) | 18.5 (0) | 201.1 (0) | 18.5 (0) | 92.96 (0) | 19.5 (0) | 53.62 (0.798) | 20 (0) |
| 18 | 87.07 | 4 | 128.78 (0.286) | 21.5 (0) | 141.58 (0.18) | 22 (0) | 42.14 (0.076) | 21 (0) | 14.18 (0) | 21.5 (0) |
| 19 | 87.99 | 2 | 80.88 (0.722) | 5 (0) | 33.86 (0.006) | 5 (0) | 1.38 (0) | 5.5 (0) | 0.48 (0) | 5.5 (0) |
| 20 | 56.97 | 5 | 128.63 (0) | 23.5 (0) | 118.11 (0) | 23 (0) | 20.76 (0) | 23.5 (0) | 11.07 (0) | 23 (0) |
| 21 | 92.31 | 3 | 89.23 (0.918) | 8 (0) | 93.12 (0.98) | 8 (0) | 39.77 (0) | 7.5 (0.002) | 16.72 (0) | 7.5 (0) |
| 22 | 102.49 | 2 | 154.38 (0.172) | 9 (0) | 153.74 (0.164) | 9.5 (0) | 78.27 (0.006) | 8.5 (0) | 39.84 (0) | 9 (0) |
| 23 | 143.49 | 7 | 202.1 (0.004) | 39 (0) | 205.85 (0) | 37.5 (0) | 202.01 (0.002) | 40 (0) | 198.13 (0) | 38 (0) |
| 24 | 70.58 | 5 | 107.35 (0.19) | 25 (0) | 114.11 (0.084) | 25 (0) | 49.49 (0.074) | 27 (0) | 21.49 (0) | 25 (0) |
| 25 | 59.48 | 4 | 156.08 (0) | 14 (0) | 156.08 (0) | 14 (0) | 116.6 (0) | 12.5 (0) | 89.98 (0.186) | 12 (0) |
| 26 | 77.47 | 6 | 83.45 (0.838) | 17 (0) | 95.44 (0.506) | 17 (0) | 69.66 (0.312) | 16.5 (0) | 38.48 (0) | 15 (0) |
| 27 | 51.61 | 2 | 47.84 (0.708) | 9.5 (0) | 39.82 (0.146) | 10 (0) | 6.26 (0) | 9.5 (0) | 3.51 (0) | 10 (0) |
| 28 | 98.45 | 4 | 45.65 (0.112) | 13.5 (0) | 41.11 (0.05) | 12.5 (0) | 8.72 (0) | 10.5 (0) | 5.08 (0) | 11 (0) |
| 29 | 74.68 | 3 | 158.42 (0.016) | 11 (0) | 60.71 (0.182) | 11 (0) | 12.57 (0) | 10.5 (0) | 5.56 (0) | 12 (0) |
| 30 | 104.49 | 7 | 34.66 (0) | 18.5 (0) | 34.09 (0) | 18.5 (0) | 9.65 (0) | 15 (0) | 7.2 (0) | 13.5 (0) |

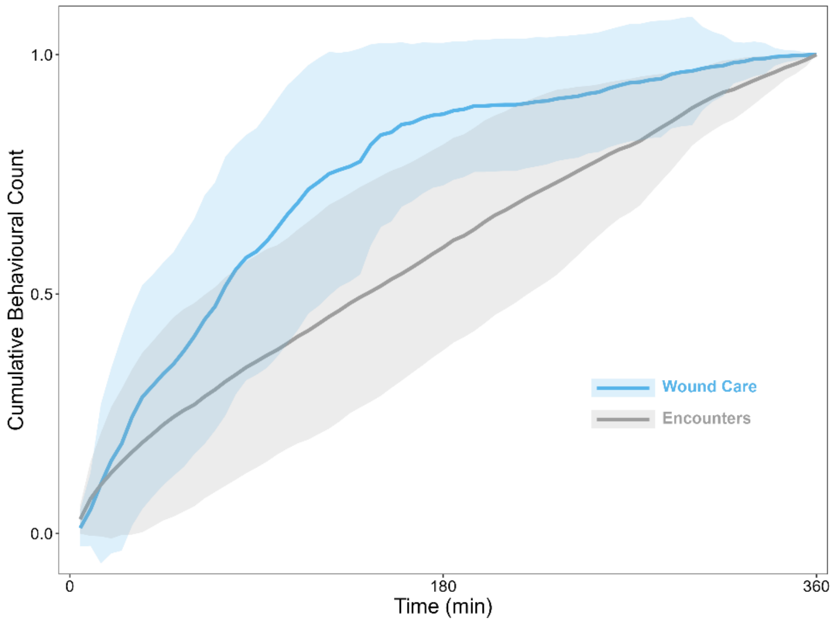


Figure S4: Cumulative distribution of wound care events received by injured ants and encounters of nestmates with injured ants during the six hours after the injury. The thicker lines represent the average cumulative distribution obtained from all injured ants (with sterile and infected wounds). The shaded region represents the standard deviation around this mean with the colors linked to the type of event in blue for wound care and grey for encounters.

#### Consistency in task performance of wound care

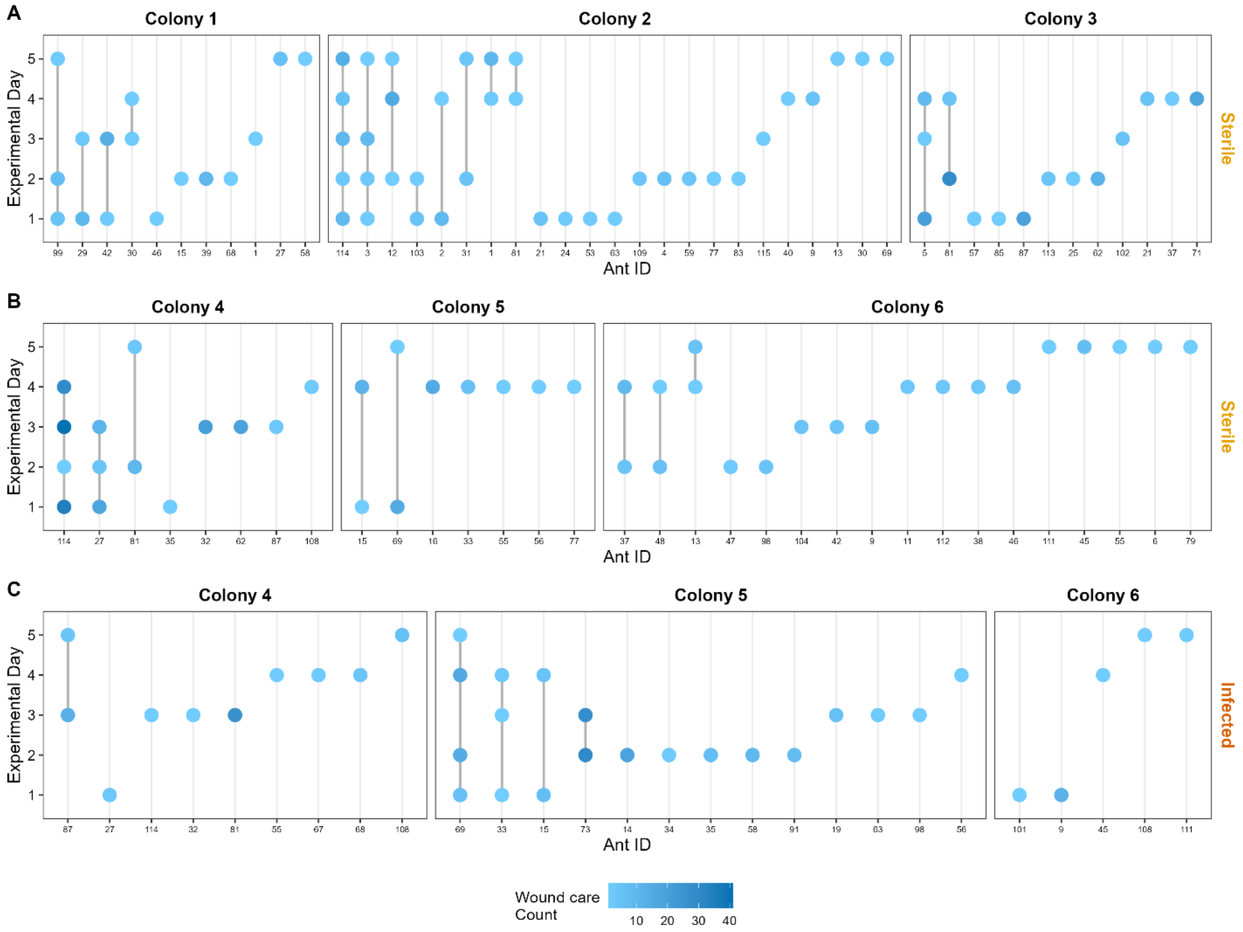


Figure S5: The number of wound care events provided by individual caregivers to ants with sterile wounds and infected wounds from all 6 colonies. Each column represents one caregiver and rows represent experimental days. Circles indicate whether the caregiver provided wound care on that day and the color represents the number of wound care events it performed. Thicker vertical lines connect circles in caregivers which performed care on multiple days.

#### Caregiver characteristics

Caregivers showed significant differences in their average displacement, proportion of time spent in the nest and nest space use entropy compared to nurses and foragers in the Pre-experimental phase (Table S10). They also had higher betweenness and degree than nurses and higher strength than foragers in this phase. Finally, they had higher space use overlap, encounters and interactions with the injured ants compared to nurses, but not foragers. Compared to non-caregivers, caregivers had higher average displacement but did not differ in the proportion of time they spent in the nest or in their nest space use entropy in the Pre-experimental phase. They also had higher betweenness and degree but not strength than non-caregivers and showed higher space use overlap and encounters with injured ants. On days in which they provided care to injured ants caregivers showed significantly higher space use overlap with injured ants in the Pre-experimental phase compared to days in which they did not provide care. Caregivers did not differ in any of their other properties on days in which they provided care compared to days in which they did not.

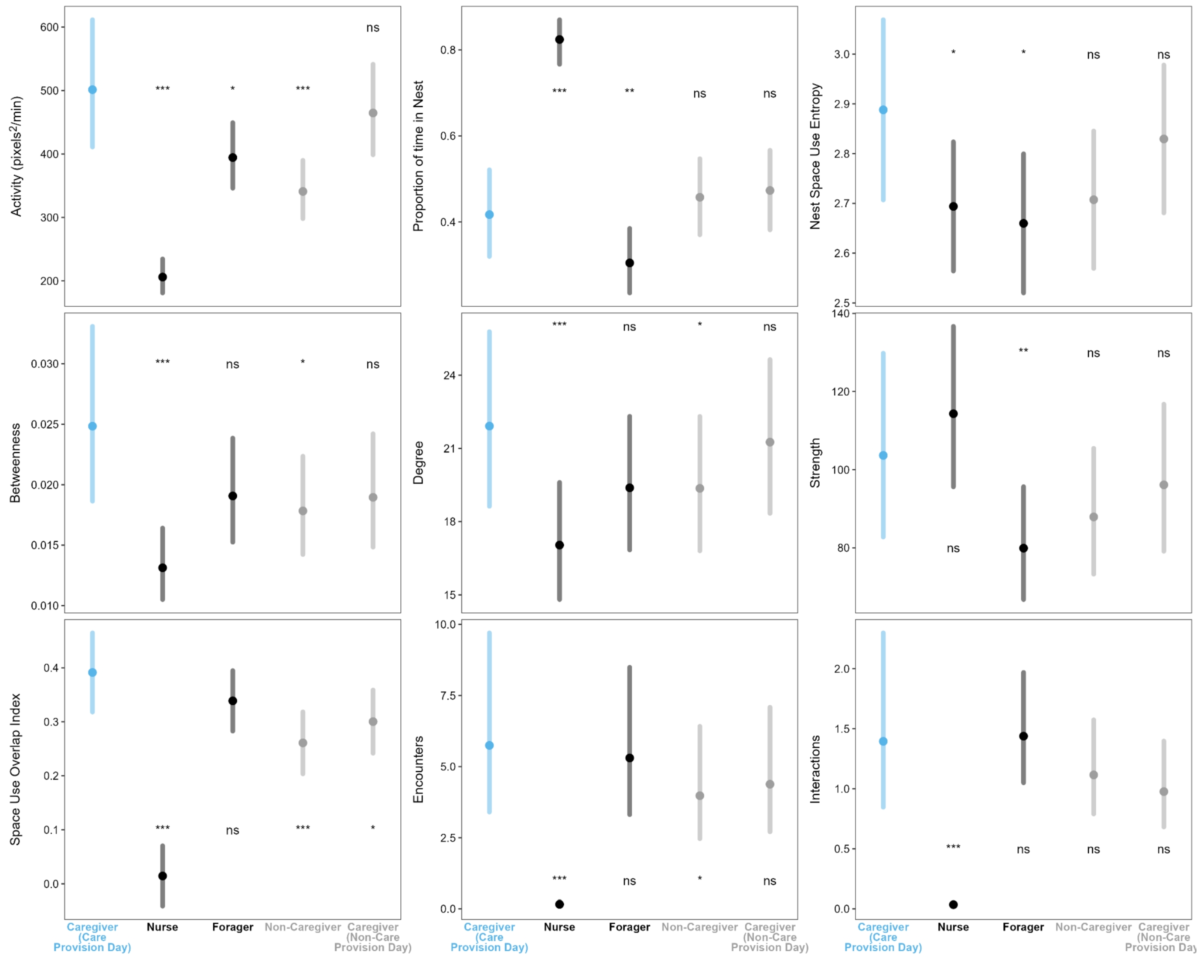


Figure S6: Differences between caregiver properties compared to other groups in the colony. Solid circles and vertical lines represent the marginal means and the 95% confidence intervals and the color represents blue for caregivers, black for the two task groups (nurse and foragers) and grey for the other two groups (non-caregivers and caregivers on days in which they did not provide care). The significance indicators for the comparison of each group with the caregivers is provided as text labels with ns for p > 0.05, * for p < 0.05, ** for p < 0.01, *** for p < 0.001.

Table S11: Estimated marginal means and confidence intervals (CI) of properties of caregivers with other groups in the Pre-experimental phase as well as the contrast difference, the t/z value and the p value associated with the difference.

| Property | Caregiver | | Comparison | | | Difference | t/z value | p value |
|---|---|---|---|---|---|---|---|---|
| | Estimate | CI | Group | Estimate | CI | | | |
| Average displacement | 501.33 | 411.04 - 611.45 | Nurse | 206.03 | 181.03 - 234.49 | -0.89 | -9.41 | < 0.001 |

| | | | | | | | | |
|---|---|---|---|---|---|---|---|---|
| | | | Forager | 394.34 | 346.11 - 449.3 | -0.24 | -2.52 | 0.042 |
| | | | Non-caregiver | 340.94 | 298.16 - 389.85 | -0.39 | -4.02 | < 0.001 |
| | | | Non care provision day | 464.61 | 398.81 - 541.26 | -0.08 | -0.72 | 0.840 |
| Proportion of time in nest | 0.42 | 0.32 - 0.52 | Nurse | 0.82 | 0.77 - 0.87 | -0.19 | -2.46 | 0.049 |
| | | | Forager | 0.30 | 0.24 - 0.39 | -0.23 | -2.69 | 0.026 |
| | | | Non-caregiver | 0.46 | 0.37 - 0.55 | -0.18 | -2.20 | 0.094 |
| | | | Non care provision day | 0.47 | 0.38 - 0.57 | -0.06 | -0.66 | 0.872 |
| Nest space use entropy | 2.89 | 2.71 - 3.07 | Nurse | 2.69 | 2.56 - 2.82 | 1.88 | 13.49 | < 0.001 |
| | | | Forager | 2.66 | 2.52 - 2.8 | -0.49 | -3.58 | 0.001 |
| | | | Non-caregiver | 2.71 | 2.57 - 2.85 | 0.16 | 1.19 | 0.562 |
| | | | Non care provision day | 2.83 | 2.68 - 2.98 | 0.23 | 1.51 | 0.366 |
| Betweenness | 0.02 | 0.02 - 0.03 | Nurse | 0.01 | 0.01 - 0.02 | -0.64 | -5.76 | < 0.001 |
| | | | Forager | 0.02 | 0.02 - 0.02 | -0.26 | -2.38 | 0.061 |
| | | | Non-caregiver | 0.02 | 0.01 - 0.02 | -0.33 | -2.97 | 0.011 |
| | | | Non care provision day | 0.02 | 0.01 - 0.02 | -0.27 | -2.22 | 0.090 |
| Degree | 21.91 | 18.63 - 25.78 | Nurse | 17.04 | 14.81 - 19.61 | -0.25 | -5.05 | < 0.001 |

| | | | | | | | | |
|---|---|---|---|---|---|---|---|---|
| | | | Forager | 19.39 | 16.84 - 22.31 | -0.12 | -2.45 | 0.051 |
| | | | Non-caregiver | 19.36 | 16.81 - 22.31 | -0.12 | -2.46 | 0.050 |
| | | | Non care provision day | 21.26 | 18.34 - 24.64 | -0.03 | -0.55 | 0.913 |
| Strength | 103.65 | 82.78 - 129.79 | Nurse | 114.32 | 95.62 - 136.68 | 0.10 | 1.19 | 0.562 |
| | | | Forager | 79.91 | 66.76 - 95.64 | -0.26 | -3.13 | 0.007 |
| | | | Non-caregiver | 87.91 | 73.27 - 105.47 | -0.16 | -1.97 | 0.155 |
| | | | Non care provision day | 96.12 | 79.13 - 116.75 | -0.08 | -0.83 | 0.787 |
| Space use overlap with injured ants | 0.39 | 0.32 - 0.46 | Nurse | 0.01 | -0.04 - 0.07 | -0.38 | -12.54 | < 0.001 |
| | | | Forager | 0.34 | 0.28 - 0.39 | -0.05 | -1.74 | 0.249 |
| | | | Non-caregiver | 0.26 | 0.2 - 0.32 | -0.13 | -4.18 | < 0.001 |
| | | | Non care provision day | 0.30 | 0.24 - 0.36 | -0.09 | -2.82 | 0.018 |
| Encounters with injured ants | 5.75 | 3.4 - 9.7 | Nurse | 0.16 | 0.1 - 0.26 | -3.59 | -24.07 | < 0.001 |
| | | | Forager | 5.30 | 3.31 - 8.49 | -0.08 | -0.60 | 0.894 |
| | | | Non-caregiver | 3.98 | 2.47 - 6.41 | -0.37 | -2.68 | 0.027 |
| | | | Non care provision day | 4.38 | 2.71 - 7.08 | -0.27 | -1.90 | 0.182 |
| Interactions with injured ants | 1.40 | 0.85 - 2.3 | Nurse | 0.04 | 0.02 - 0.05 | -3.68 | -14.00 | < 0.001 |

| | | | | | | | | |
|---|---|---|---|---|---|---|---|---|
| | | | Forager | 1.44 | 1.05 - 1.97 | 0.03 | 0.14 | 0.997 |
| | | | Non-caregiver | 1.12 | 0.79 - 1.57 | -0.22 | -0.97 | 0.704 |
| | | | Non care provision day | 0.98 | 0.68 - 1.4 | -0.36 | -1.47 | 0.387 |

#### Prediction of care behavior

Loadings for each principal component of each property can be found in Table S11. M1 positively correlated with more active individuals, M2 negatively correlated with individuals which covered the nest more evenly, N1 with more interactive individuals, N2 with more central individuals but with lower average number of interactions, A1 with individuals who had greater affinity with the injured ants and A2 with individuals less connected to injured ants in the interaction network. For N1 and A1, the original component correlated negatively with the property with the highest weights, we therefore inverted the signs of both components to make interpretation easier.

Table S12: Loadings of the different principal components of each property used to predict care behavior.

| Property Type | Property | Principal Component | |
|---|---|---|---|
| | | 1 | 2 |
| Movement | Total displacement | 0.60 | -0.31 |
| | Average displacement | 0.61 | 0.30 |
| | Nest space use entropy | -0.16 | -0.75 |
| | Proportion of time in Nest | -0.50 | -0.50 |
| Network | Betweenness | 0.31 | -0.61 |
| | Degree | 0.53 | -0.37 |
| | Strength | 0.61 | 0.16 |
| | Average number of interactions | 0.47 | 0.45 |
| | Social maturity | -0.14 | -0.51 |

| | | | |
|---|---|---|---|
| Affinity with injured ants | Space use overlap | 0.57 | 0.05 |
| | Number of encounters | 0.59 | 0.08 |
| | Number of interactions | 0.52 | 0.26 |
| | Distance in the interaction network | -0.22 | 0.96 |

Model comparisons using BIC revealed that models which included A1 and A2 were the best supported by the data (Table S12). Models which included single predictors like the encounters or interactions with injured ants in the Pre-experimental phase had poor support compared to the models with A1 and A2. We also tested whether A1 and A2 predicted the amount of wound care provided and found that A1 had a significant effect on the amount of wound care provided in addition to its effect on the probability of care provisioning (Table S13). Furthermore, A1 and A2 showed the same qualitative pattern even when we only considered ants which interacted at least once with the injured ants in the Experimental phase (i.e., only caregivers and non-caregivers, Table S13).

Table S13: The 12 different models used for the model comparison along with the predictors in the model the BIC, ΔBIC and relative weight associated with each model. Model numbers correspond to those in Table S2.

| Model | Predictors from Pre-experimental phase | Bayesian Information Criterion | | |
|---|---|---|---|---|
| | | BIC | ΔBIC | Weight |
| M41 | A1 + A2 | 1090.66 | 0.00 | 0.903 |
| M44 | Overlap index | 1096.27 | 5.61 | 0.055 |
| M38 | Overlap index + Encounters + Interactions + Network distance | 1097.34 | 6.68 | 0.032 |
| M35 | M1 + M2 + N1 + N2 + A1 + A2 | 1100.55 | 9.89 | 0.006 |
| M37 | Betweenness + Degree + Strength + Average number of interactions + Social maturity | 1102.38 | 11.72 | 0.003 |
| M43 | Encounters | 1103.79 | 13.13 | 0.001 |
| M45 | Social Maturity | 1106.33 | 15.67 | < 0.001 |

| | | | | |
|---|---|---|---|---|
| M34 | Average displacement + Total displacement + Nest space use entropy + Proportion of time in Nest + Betweenness + Degree + Strength + Average number of interactions + Overlap index + Encounters + Interactions | 1111.67 | 21.01 | < 0.001 |
| M39 | M1 + M2 | 1113.08 | 22.42 | < 0.001 |
| M40 | N1 + N2 | 1118.64 | 27.98 | < 0.001 |
| M36 | Average displacement + Total displacement + Nest space use entropy | 1138.14 | 47.48 | < 0.001 |
| M42 | Interactions | 1148.67 | 58.01 | < 0.001 |

Table S14: Odds ratio of the probability of providing care and incidence rate ratio of the amount of care along with the confidence intervals and the p value for A1 and A2 which best predicted the probability of care provision. The results for both sets of comparison – one with all ants and another with only data from caregivers and non-caregivers is provided.

| Comparison | Predictor | Probability of care | | | Amount of care | | |
|---|---|---|---|---|---|---|---|
| | | Odds ratio | CI | p value | Incidence rate ratio | CI | p value |
| All ants | Intercept | 0.03 | 0.02 - 0.05 | < 0.001 | 0.20 | 0.15 - 0.27 | < 0.001 |
| | A1 | 1.54 | 1.41 - 1.69 | < 0.001 | 1.64 | 1.26 - 2.13 | <0.001 |
| | A2 | 0.76 | 0.64 - 0.91 | 0.003 | 0.79 | 0.53 - 1.17 | 0.236 |
| Caregivers and non-caregivers | Intercept | 0.16 | 0.12 - 0.20 | < 0.001 | 0.89 | 0.61 - 1.28 | 0.523 |
| | A1 | 1.31 | 1.17 - 1.46 | < 0.001 | 1.18 | 1.00 - 1.39 | 0.046 |
| | A2 | 0.76 | 0.61 - 0.95 | 0.015 | 0.78 | 0.55 - 1.12 | 0.176 |

698
